# HIV-1 initiates genomic RNA packaging in a unique subset of host RNA granules

**DOI:** 10.1101/183855

**Authors:** Motoko Tanaka, Brook C. Barajas, Bridget A. Robinson, Daryl Phuong, Kasana Chutiraka, Jonathan C. Reed, Jaisri R. Lingappa

## Abstract

How HIV-1 genomic RNA (gRNA) is packaged into assembling virus remains unclear. Here, we use biochemical and *in situ* approaches to identify the complex in which the capsid protein Gag first associates with gRNA, termed the packaging initiation complex. First, we show that in the absence of assembling Gag, non-nuclear non-translating gRNA is nearly absent from the soluble fraction of provirus-expressing cells, and is found instead primarily in complexes >30S. When we express a Gag mutant known to be arrested at packaging initiation, we find only one complex containing Gag and gRNA; thus, this complex corresponds to the packaging initiation complex. This ∼80S complex also contains two cellular facilitators of assembly, ABCE1 and the RNA granule protein DDX6, and therefore corresponds to a co-opted host RNA granule and a previously described capsid assembly intermediate. Additionally, we find this granule-derived packaging initiation complex in HIV-1-infected H9 T cells, and demonstrate that wild-type Gag forms both the packaging initiation complex and a larger granule-derived complex corresponding to a late packaging/assembly intermediate. We also demonstrate that packaging initiation complexes are far more numerous than P bodies *in situ*. Finally, we show that Gag enters the ∼80S granule to form the packaging initiation complex via a two-step mechanism. In a step that is independent of a gRNA-binding domain, Gag enters a broad class of RNA granules, most of which lack gRNA. In a second step that is dependent on the gRNA-binding nucleocapsid domain of Gag or a heterologous gRNA-binding domain, Gag enters a gRNA-containing subset of these granules. Thus, we conclude that packaging in cells does not result from random encounters between Gag and gRNA; instead our data support a fundamentally different model in which Gag is directed to gRNA within a unique host RNA granule to initiate this critical event in HIV-1 replication.

**Nontechnical Summary:** To form infectious virus, the HIV-1 capsid protein Gag must associate with and package the viral genomic RNA (gRNA) during the virus assembly process. HIV-1 Gag first associates with gRNA in the cytoplasm, forming a complex termed the packaging initiation complex; this complex subsequently targets to the plasma membrane where Gag completes the assembly and packaging process before releasing the virus from the cell. Although the packaging initiation complex is critical for infectious virus formation, its identity and composition, and the mechanism by which it is formed, remain unknown. Here we identify the packaging initiation complex, and demonstrate that it corresponds to a host RNA granule that is co-opted by the virus. RNA granules are diverse complexes utilized by host cells for all aspects of RNA storage and metabolism besides translation. Our study also defines the mechanism by which HIV-1 Gag enters this host RNA granule to form the packaging initiation complex, and reveal that it involves two steps that depend on different regions of Gag. Our finding that Gag co-opts a poorly studied host complex to first associate with gRNA during packaging provides a new paradigm for understanding this critical event in the viral life cycle.

---

**Abbreviations used**
αABCEl: antibodies directed against ABCE1
αDDX6: antibodies directed against DDX6
αGag: antibodies directed against Gag
αGFP: antibodies directed against GFP
coIP: coimmunoprecipitation
gRNA: HIV-1 genomic RNA
IEM: immunoelectron microscopy
IP: immunoprecipitation
LZ: leucine zipper
NC: nucleocapsid
O-FISH: oligo fluorescent in situ hybridization
O-FISH PLA: oligo fluorescent in situ hybridization coupled with the proximity ligation assay PLA, proximity ligation assay PM, plasma membrane
PuroHS: treatment with puromycin plus high salt
RNPC: ribonucleoprotein
RT-qPCR: 
SRP: signal recognition particle
VLPs: virus-like particles
WB: Western blot
WT: wild type

## Introduction

For released HIV-1 particles to be infectious, two copies of full-length genomic RNA (gRNA) must be packaged during assembly of the immature HIV-1 capsid. Capsid assembly involves oligomerization of Gag in the cytoplasm followed by targeting of Gag via its N-terminal myristate to the plasma membrane (PM), where ∼3000 Gag proteins multimerize to form each immature capsid. Packaging of gRNA is initiated when the structural protein Gag first associates with gRNA during assembly, and requires the nucleocapsid domain (NC) of Gag as well as specific encapsidation signals in gRNA (reviewed in [1]). The gRNA-containing immature capsids subsequently undergo budding, release, and maturation (reviewed in [2]). In addition to being used for packaging, HIV-1 gRNA in the cytoplasm is also used for translation of Gag and GagPol (reviewed in [1]). Translation and packaging are thought to be mutually exclusive, with the shift between these processes likely controlled by a gRNA conformational switch (reviewed in [3,4]). Thus, gRNAs that are packaged are likely to be nontranslating gRNAs with unique structural features [5]. In the absence of gRNA, capsid assembly and release still occur but the resulting virus is non-infectious [6].

The process by which HIV-1 Gag finds gRNA at the start of packaging is of great interest. Biochemical and imaging studies have established that a Gag dimer or oligomer first associates with gRNA in the cytoplasm, forming a complex termed the packaging initiation complex [7]. Subsequently, the packaging initiation complex targets to the PM where Gag multimerizes to form the immature capsid [8]. Live imaging studies have defined the diffusion characteristics of Gag and gRNA in cells [9,10]; but, notably, the exact components in these diffusing complexes have not been identified. Thus, although formation of the packaging initiation complex is critical for production of infectious virus, the composition of the packaging initiation complex and the mechanism by which Gag and gRNA first associate to form this complex are not known.

Here we used our understanding of HIV-1 capsid assembly to determine how the packaging initiation complex is formed. Our previous studies showed that assembling HIV-1 Gag co-opts RNA granules [11], which are host ribonucleoprotein complexes (RNPCs) that contain non-translating mRNAs. While all translating cellular mRNAs are in monosomes or polysomes, non-translating cellular mRNAs reside in RNA granules (reviewed in [12]). Numerous types of RNA granules exist; these granules are distinguished by sizes and marker proteins, and function in silencing, storage, degradation, stress, and other events in RNA metabolism. Some RNA granules, such as P bodies and stress granules, are easily visible by light microscopy, but others are smaller and poorly understood. We found that after co-opting host RNA granules, Gag remains in these granules during immature capsid assembly, forming a sequential pathway of assembly intermediates named by their sedimentation values of ∼80S, ∼150S, and ∼500S (reviewed in [13]). Both the original co-opted host RNA granule and the subsequently formed assembly intermediates contain multiple RNA granule proteins, including the DEAD-box RNA helicase DDX6 [11].

These previous studies led us to what we term the RNA granule model of packaging. In this model, we propose that, like non-translating cellular mRNA, non-translating gRNAs are sequestered within RNA granules; thus, Gag must enter gRNA-containing host RNA granules to initiate gRNA packaging. Localizing packaging to RNA granules could be advantageous to the virus in a number of ways: it would sequester gRNA from the host innate immune system, concentrate Gag at the site where gRNA is located, and place packaging and assembly in proximity with host enzymes that could facilitate those events. In keeping with the latter possibility, two of the host proteins present in both the assembly intermediates and the host RNA granules from which they are derived are cellular facilitators of HIV-1 capsid assembly – the ATP-binding cassette protein E1 (ABCE1) and DDX6 [11,14].

Here we identified the packaging initiation complex and tested the RNA granule model of HIV-1 packaging. First we showed that all non-translating HIV-1 gRNAs are in large complexes, even in the absence of assembling Gag. Using a Gag mutant that is arrested after packaging initiation, we demonstrated that only one complex contains Gag associated with gRNA, and therefore fits the definition of the packaging initiation complex. This complex is an ∼80S RNA granule that also contains the cellular facilitators of assembly, ABCE1 and DDX6. *In situ* studies confirmed these findings and revealed that the packaging initiation complexes are smaller and more numerous than P bodies. Additionally, we examined the mechanism by which the packaging initiation complex is formed, and found that Gag uses both a gRNA-binding-independent and a gRNA-binding-dependent step to localize to gRNA-containing RNA granules. Finally, we confirmed the physiological relevance of the RNA-granule derived packaging intermediates by identifying them in a chronically infected human T cell line. Together, our data argue that packaging of the HIV-1 genome is initiated only after Gag localizes to a unique and poorly understood subclass of host RNA granules that contains non-translating gRNA.

## Results

### Confirmation of packaging phenotypes for proviruses expressing WT Gag and Gag mutants

To study gRNA packaging, we used a variety of gRNA expression systems (Fig 1A) that produce either WT Gag or Gag mutants with known phenotypes (Fig 1B). WT Gag forms the packaging initiation complex and completes assembly to produce virus-like particles (VLPs) that contain gRNA. Two assembly-defective Gag mutants, MACA Gag and G2A Gag, were studied because they arrest Gag assembly before or after packaging initiation complex formation, respectively. The assembly-incompetent MACA Gag fails to form the packaging initiation complex because it lacks the gRNA-binding NC domain (reviewed in [13,15,16]). In contrast, the assembly-defective G2A Gag forms the cytoplasmic packaging initiation complex [7] but is arrested in the cytoplasm due to a point mutation that prevents the myristoylation required for PM targeting [17–19]. We also analyzed the assembly-competent but gRNA-interaction-deficient HIV-1 GagZip chimera. In place of the gRNA-binding NC domain, GagZip contains a dimerizing leucine zipper (LZ), which supports assembly but not RNA association [20–24]; thus GagZip produces VLPs that lack gRNA, and would not be expected to form the packaging initiation complex. GagZip is of interest since a packaging model should be able to explain how GagZip successfully assembles capsids but fails to incorporate gRNA.

**Fig 1.**
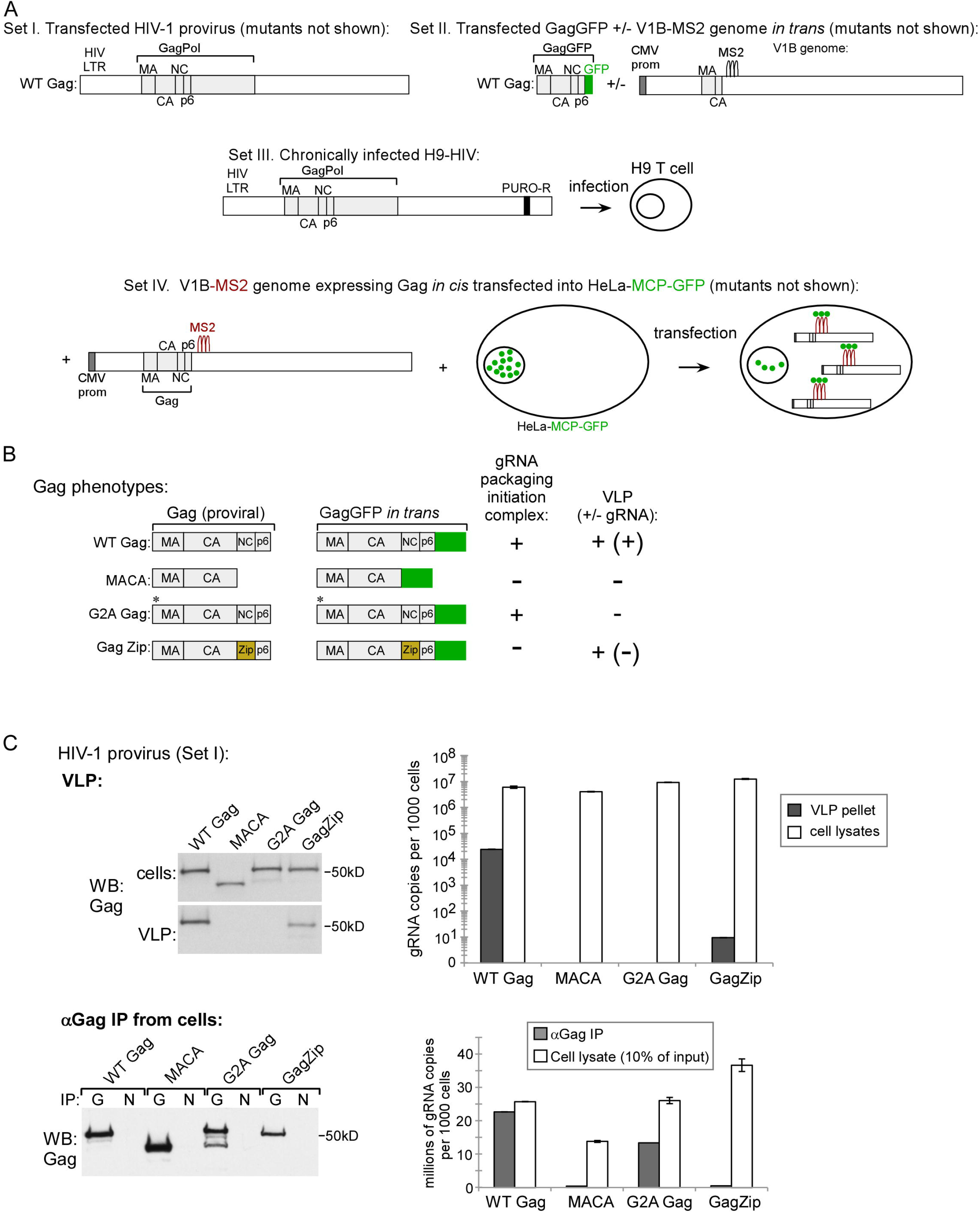
Gag constructs: diagrams and phenotyopes. **A)** Diagram of the different *cis* and *trans* expression systems used (Sets I - IV). Only WT constructs are shown here, with mutant constructs diagrammed in later figures. **Set I** consists of WT (and mutant) HIV-1 proviruses (pro-, delta env). **Set II** consists of codon-optimized WT (and mutant) Gag constructs fused to GFP (GagGFP), transfected with a modified genomic construct (V1B) provided *in trans.* Set III consists of proviruses from Set I in which the *nef* gene was replaced with a puromycin resistance gene (PURO-R). H9 T cells were infected with these constructs and maintained under puromycin selection to generate a chronically infected H9 T cell line expressing WT Gag (H9-HIV). In **Set IV** consists of V1B constructs (see Set II), which were engineered to express WT or mutant Gag *in cis.* Set VI constructs were transfected into HeLa-MCP-GFP cells, which express a GFP-tagged MS2 capsid protein (MCP) that contains a nuclear localization signal. Since V1B genomic constructs also contain MS2 binding sites, MCP-GFP binds to V1B, resulting in GFP tagging of V1B gRNA [8], which allows subsequent immunolabeling of V1B gRNA with GFP antibody. **B) Summary of expected Gag phenotypes**. Diagram indicates, for WT Gag and Gag mutants, whether the packaging initiation complex is predicted to form and whether VLPs are known to be produced. Released VLPs that contain or lack gRNA are indicated by +(+) and +(-) respectively. **C) Confirmation of expected phenotypes for VLP production and packaging initiation complex formation**. COS-1 cells were transfected with Set I constructs from A. Top row: WB shows Gag in cell lysates and VLPs. Graph shows gRNA copy number, as determined by RT-qPCR, in VLPs or cell lysates representing the equivalent of 1000 cells. Similar results were obtained in 293T cells. Bottom row: Cell lysates expressing proviruses encoding WT or mutant Gag (Set I constructs in panel A) were subjected to IP with αGag or nonimmune (N) antibody followed by Gag WB (left). IP eluates were also analyzed by RT-qPCR for gRNA copies, with NI values subtracted (graph). Graph shows that only WT and G2A Gag form intracellular complexes containing Gag associated with gRNA, thereby confirming packaging initiation complex phenotypes shown in B. Error bars show SEM from duplicate samples. Data are representative of three independent repeats.

We confirmed the VLP phenotypes described above following transfection of WT and mutant proviruses (Fig 1A, Set I) into COS-1 (Fig 1C) by analyzing cell lysates and VLPs for Gag and gRNA copies (Fig 1C, top). Additionally, the predicted packaging initiation complex phenotypes of these four Gag constructs (Fig 1B) were confirmed by quantifying gRNA copies associated with intracellular Gag, using immunoprecipitation (IP) from cell lysates with antibody directed against Gag (αGag), followed by RTqPCR of IP eluates (Fig 1C, bottom).

### Non-translating gRNAs are in diverse large complexes even in the absence of assembling Gag

To date, the spectrum of gRNA-containing complexes present in the cytoplasm has not been defined. Our model predicts that gRNA behaves like mRNA; thus, even in the absence of Gag we would expect to find that all HIV-1 gRNA is either in translating complexes or in diverse non-translating RNPCs. Moreover, because gRNA, like mRNA, should associate with ribonucleoproteins to form relatively large complexes, we would not expect to find gRNA in the soluble fraction of cell lysates, which contains monomers, dimers, and small oligomers. To test these predictions, we transfected cells with MACA provirus (Fig 2A), which expresses an otherwise full-length gRNA encoding a truncated assembly-incompetent MACA Gag protein that is arrested as an ∼10S complex [25]. These cells were harvested under conditions that removed nuclei, kept RNPCs intact, and solubilized any membranes associated with proteins. Lysates were analyzed by velocity sedimentation followed by RT-qPCR to determine the approximate size (S value) of all non-nuclear RNPCs containing gRNA (or other RNAs). This analysis was performed with both translating and non-translating complexes intact, and also following disruption of translating ribosomes with *puromycin* and high salt treatment (PuroHS). PuroHS releases nascent chains from ribosomes, dissociates ribosomal subunits, and releases mRNA [26,27], which then likely shifts into non-translating RNPCs [28]. Thus, PuroHS treatment should allow analysis of only non-translating gRNAs, the population likely to undergo packaging.

**Fig 2:**
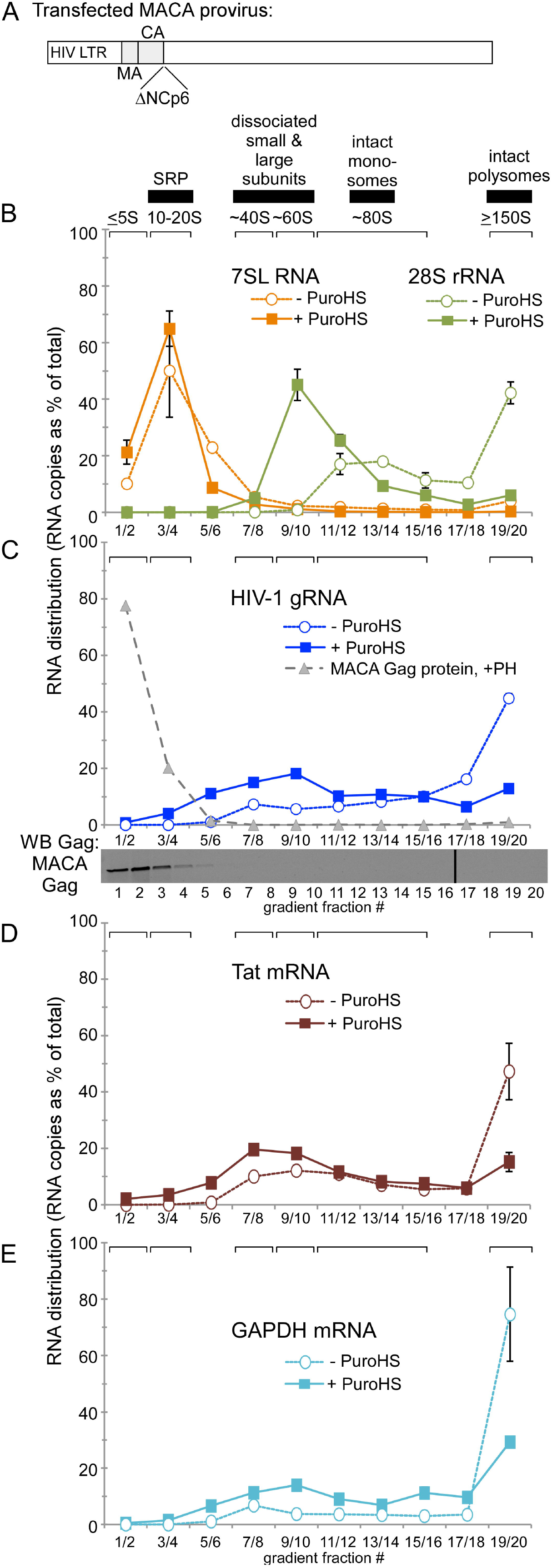
Non-translating HIV-1 gRNA is primarily in diverse complexes >30S in the absence of assembling Gag. **A-E)** Lysate from COS-1 cells transfected with the assembly-incompetent MACA provirus (Fig 1A, Set I) was divided into two pools, that were either not treated or treated with PuroHS (-/+ PuroHS). Both pools were analyzed in parallel by velocity sedimentation followed by RTqPCR of paired gradient fractions using the appropriate qPCR primer sets to determine copy number of the indicated RNAs (28SrRNA, 7SL RNA, HIV-1 gRNA, HIV-1 Tat mRNA, or GAPDH mRNA). Quantity of the indicated RNAs in gradient fractions is expressed as a distribution (% of RNA in all fractions) to allow comparison between different RNAs. Blot in panel C shows migration of MACA protein, which is also graphed as a gray dotted line in C. Brackets at top show S value markers, and horizontal bars show expected migrations of various RNPCs. Error bars show SEM from duplicate samples. Data are from a single experiment that is representative of three independent repeats.

We first determined the efficacy of ribosome disruption by PuroHS. To do this, we analyzed MACA gradient fractions for 28S rRNA, a marker for the 60S large ribosomal subunit (Fig 2B), following PuroHS treatment or in the absence of PuroHS treatment. As expected, in the absence of PuroHS treatment (Fig 2B, open green circles), 28S rRNA migrated almost entirely in the position of monosomes (∼80S) and polysomes (>150S). Following PuroHS treatment (Fig 2B, solid green squares), 28S rRNA migrated almost entirely in an ∼60S peak representing the dissociated large ribosomal subunit, with almost no 28S rRNA remaining in the polysome region (>150S). The near complete absence of 28S rRNA in the polysome region after PuroHS treatment indicated highly effective ribosome disassembly. From these data we concluded that PuroHS disassembles most 28S-containing monosomes and polysomes into large and small ribosomal subunits. As a negative control, the same fractions were examined for 7SL RNA, a marker for signal recognition particle (SRP), which is a ribosome-independent ∼11S RNPC [29]. As expected, PuroHS treatment did not affect the migration of 7SL (Fig 2B, compare open orange circles vs. solid orange squares). Thus, RNPCs of different sizes (monosomes, polysomes, and SRP) remain intact following cell lysis and velocity sedimentation, and PuroHS treatment disrupted translating RNPCs, but not ribosome-independent RNPCs.

Analysis of HIV-1 gRNA in these same MACA fractions revealed that, with or without PuroHS treatment, almost no HIV-1 gRNA was in the soluble fraction (< 20S) and very little was in small complexes of < 30S (Fig 2C, compare open blue circles to solid blue squares). Thus, at steady state in the absence of assembling Gag, most gRNA is in complexes > 30S, consisting of translating complexes and non-translating complexes. This contrasted with MACA protein, which was found almost entirely in the soluble fraction (< 20S; Fig 2C, grey triangles). Following PuroHS treatment, > 95% of gRNA was in diverse RNPCs > 30S that formed a broad peak centered at ∼60S. Comparison of gRNA data obtained with and without PuroHS treatment suggests that approximately 30% of total gRNA was in translating polysomes, defined as complexes > 150S that are lost upon PuroHS. Moreover, PuroHS treatment caused HIV-1 gRNA that was in polysomes (>150S) to shift into non-translating RNPCs of ∼30-150S, some of which are present even in the absence of PuroHS treatment (Fig. 2C). When the same gradient fractions were analyzed for HIV-1 subgenomic Tat mRNA and a cellular mRNA (GAPDH), a pattern similar to that of HIV-1 gRNA was observed, with or without PuroHS treatment (compare Fig 2D, E to 2C), although perhaps with GAPDH mRNA forming more >150S complexes after puromycin treatment, which could correspond to non-translating RNA granules or residual polysomes not disrupted by puromycin. Thus, the distribution of translating and non-translating gRNA-containing complexes resembles the distribution of cellular mRNAs and subgenomic viral mRNAs. Because packaging likely involves non-translating gRNA, as explained above, all subsequent lysate experiments utilized PuroHS to allow analysis of almost exclusively non-translating gRNA.

### G2A Gag enters gRNA-containing granules to form the ∼80S packaging initiation complex

Having shown that non-translating gRNA is primarily in RNPCs > 30S, and does not comigrate with soluble packaging-incompetent MACA Gag, we next asked whether a packaging-initiation-competent Gag enters one of the > 30S RNPCs to form the packaging initiation complex. Because the packaging initiation complex is the complex in which Gag and gRNA first associate, we would expect the packaging initiation complex to have two key features – first, gRNA and Gag should comigrate in the fractions that contain the packaging initiation complex, and secondly, gRNA and Gag in these fractions should be in association with each other, through direct or indirect interactions, as indicated by coimmunoprecipitation (coIP). In addition, if the packaging initiation complex is a co-opted RNA granule, we would expect gRNA to associate with RNA granule proteins in these fractions.

To help identify the packaging initiation complex, we utilized a provirus that expresses G2A Gag. G2A Gag forms the cytoplasmic complex that has been defined as the packaging initiation complex [7], and is likely arrested in the form of the packaging initiation complex due to its inability to target to membranes. To demonstrate comigration of gRNA with packaging-initiation-competent G2A Gag, cells expressing MACA vs. G2A proviruses (Fig 1A, Set I) to similar steady state levels (Fig 3A), were analyzed by velocity sedimentation followed by quantitation of gRNA in gradient fractions (Fig 3B). Cells expressing either the MACA or G2A provirus displayed the same distribution of total non-nuclear non-translating gRNA, with gRNA mainly in ∼30-80S RNPCs in both cases (Fig 3B, graph). In contrast, the distribution of the MACA vs. G2A Gag protein across the gradient differed dramatically (Fig 3B, WB below graph), as observed previously [25,30]. While MACA protein was almost entirely in the soluble fraction (Fig 3B, MACA WB, fractions 1 - 3), G2A Gag protein was present in both the < 20S soluble fraction and in ∼40-80S complexes (Fig 3B, G2A WB, fractions 8 - 12). Thus, a population of packaging-initiation-competent G2A Gag protein comigrated with gRNA in ∼40-80S RNPCs, in contrast to the MACA protein, which is not competent for packaging initiation.

**Fig 3:**
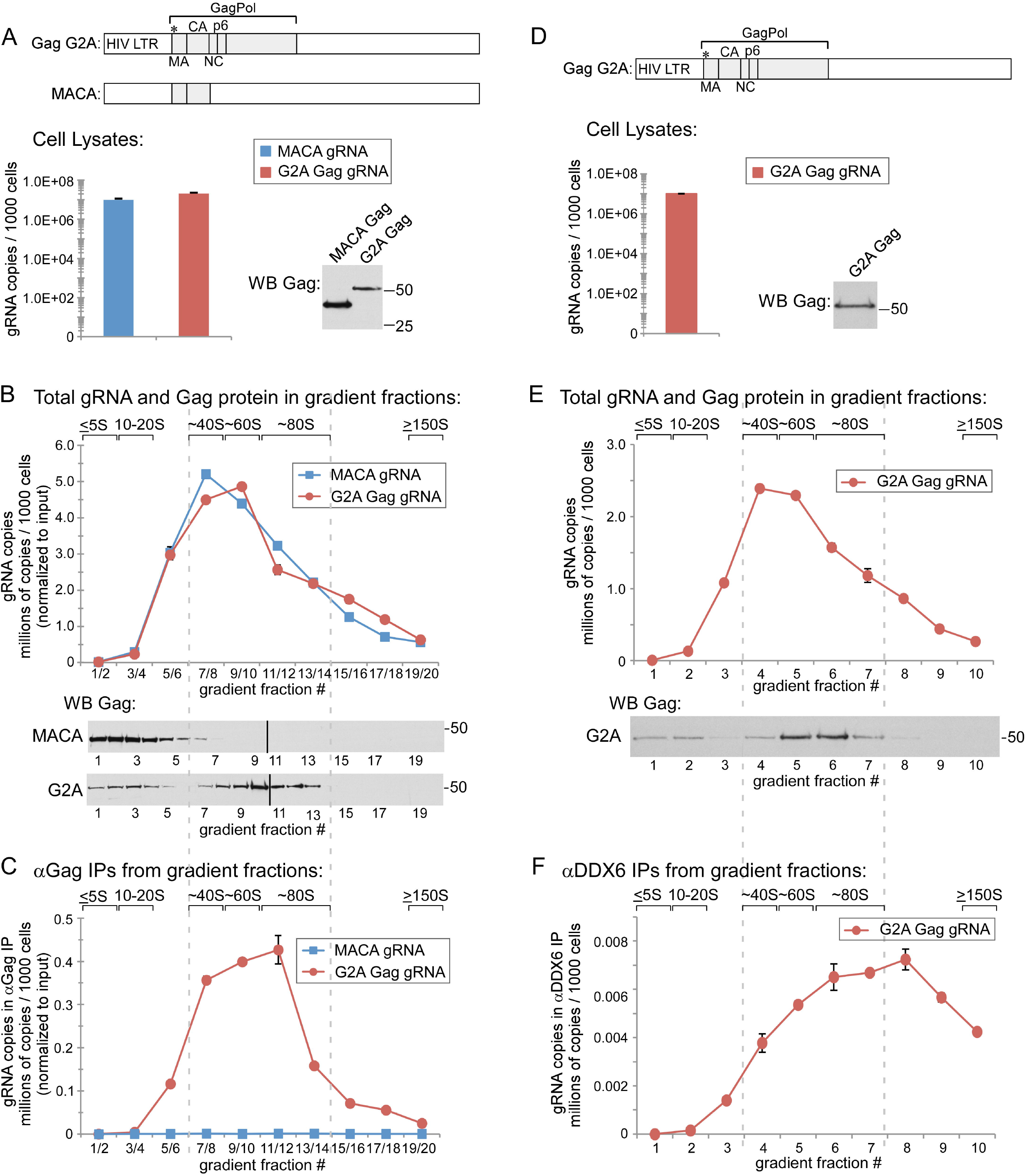
The packaging initiation complex is an ∼80S RNA granule. **A)** COS-1 cells transfected with indicated proviruses (Fig 1A, Set I) were harvested follwing PuroHS treatment, and gRNA copy number per 1000 cells was determined. **B)** Lysates from A were analyzed by velocity sedimentation, and gRNA copies per 1000 cells wasdetermined and normalized to the inputs shown in A. **C)** Gradient fractions from B were subjected to IP with HIV immune globulin (αGag), and gRNA copy number in IP eluates per 1000 cells was determined and normalized to the inputs shown in A. Similar results were obtained upon IP with monoclonal antibody to p24. **D)** COS-1 cells transfected with the indicated G2A Gag provirus (Fig 1A, Set I) were harvested following PuroHS treatment, and total gRNA copy number per 1000 cells was determined. **E)** Lysates from D were also analyzed by velocity sedimentation, and gRNA copy number per 1000 cells in each fraction was determined. **F)** Gradient fractions from E were subjected to IP with αDDX6, and IP eluates from each fraction were analyzed for gRNA copy number per 1000 cells. Brackets at top show S value markers, and dotted lines demarcate assembly intermediates based on their WB migrations. Error bars show SEM from duplicate samples. Data in each column are from a single experiment that is representative of three independent repeats.

To determine whether gRNA in the ∼40-80S RNPCs is associated with comigrating G2A Gag, gradient fractions in Fig 3B were subjected to αGag IP, followed by quantitation of gRNA in IP eluates (Fig 3C). Note that all IPs in this study were performed under native conditions, and are thus expected to pull down the protein targeted by the antibody as well as other components stably associated with the target protein through direct or indirect interactions. Even though αGag coimmunoprecipitated MACA effectively (Fig 1C), no gRNA was associated with MACA by αGag IP in any fraction, as expected given that MACA does not comigrate with gRNA (Fig 3B, compare graph to blot). Similarly, gRNA was not associated with soluble G2A Gag protein (in the < 20S region) by coIP, as expected given the lack of gRNA in the soluble fractions. Thus, the packaging initiation complex was not found in soluble fractions. In contrast, G2A Gag protein was strongly associated with gRNA by αGag coIP in a broad RNPC that peaked at ∼80S, heretofore referred to as the ∼80S complex (Fig 3C, graph). From these data, we conclude only one RNPC - the ∼80S complex - fits the definition of the packaging initiation complex in that it contains G2A Gag associated with gRNA.

Next we asked whether this ∼80S RNPC is an RNA granule, by determining if gRNA in this complex is associated with the RNA granule protein, DDX6. DDX6 is a marker for P bodies [31], but is also found in ∼80S and ∼500S RNA granules that are co-opted by Gag during assembly and contain the host enzyme ABCE1 [11]. In cells expressing G2A provirus (Fig 3D), gRNA was again observed primarily in the ∼40-80S region of gradients (Fig 3E). Moreover, antibody to DDX6 (αDDX6) coimmunoprecipitated gRNA from a complex that peaks in the ∼80S region (Fig 3F). These findings are consistent with our previous observation that DDX6 is associated with Gag in the ∼80S assembly intermediate by coIP [11]. Taken together, these data suggest that to form the packaging initiation complex, packaging-competent G2A Gag must enter an ∼80S DDX6-containing RNA granule that contains non-translating gRNA. Notably, gRNA is absent from the <20S fraction of the cell lysate, with or without PuroHS treatment (Fig 2C), and in the presence or absence of packaging-initiation-competent Gag (Fig 3B); thus we could find no evidence of a small complex containing only gRNA and a dimer of Gag.

### WT Gag forms the ∼80S packaging initiation complex and ∼500S late packaging intermediate, which contain RNA granule proteins

Previously, we used pulse-chase and mutational approaches to demonstrate that WT Gag forms an ∼80S early capsid assembly intermediate, a ∼500S late assembly intermediate, and a ∼750S complex that corresponds to a completed immature capsid [25,32–34]. Additionally, we had used coIPs to demonstrate that the ∼80S and ∼500S assembly intermediates contain the assembly facilitators ABCE1 and DDX6 in association with Gag [11,14]. If the ∼80S assembly intermediate is also the earliest packaging intermediate, as suggested by the data above (Fig 3), then we would expect the ∼500S assembly intermediate, which is only found at the PM [25], to be a late packaging intermediate that contains gRNA in association with targeting-competent Gag. Thus, WT Gag should be associated with gRNA in both the ∼80S packaging initiation complex/early assembly intermediate and the ∼500S late assembly/packaging intermediate, as well as in any intracellular ∼750S completed immature capsids that accumulate prior to virus budding and release. To test these predictions, we needed a method for immunoprecipitating multimerized Gag. Since diverse Gag antibodies fail to IP multimerized Gag because of epitope masking [35], we utilized αGFP IP of GFP-tagged, codon-optimized Gag (GagGFP) cotransfected with VIB, a modified proviral construct that expresses a gRNA containing all the signals needed for packaging [7]. Notably, V1B encodes a truncated assembly-incompetent Gag that does not interfere with assembly of GagGFP expressed in trans. These constructs coexpressed *in trans* (GagGFP/V1B, Fig 1A, Set II) have been well vetted in live imaging studies [8]. Use of this *in trans* system also allowed us to express a single well-studied V1B genomic construct with different Gag constructs. As expected, coexpression of GagGFP and V1B constructs resulted in the same VLP and packaging-initiation-complex phenotypes observed for proviral constructs in Fig 1B (Fig S1A).

To test whether WT Gag forms the ∼80S putative packaging initiation complex and larger ∼500S and ∼750S complexes, we analyzed cell lysates coexpressing either WT Gag with V1B or G2A GagGFP with the V1B genomic construct. In this experiment, we used a velocity sedimentation gradient that allows resolution of early and late assembly intermediates [25,30]. Steady state levels of intracellular Gag protein were similar for both WT and G2A, as were gRNA levels (Fig 4A). G2A Gag protein formed only the ∼10S and ∼80S intermediates (as in Fig 3B); in contrast, at steady state, WT Gag protein formed both of these complexes as well as the ∼500S late assembly intermediate and ∼750S completed capsids (Fig 4B, WB), as shown previously [25,30,32]. Notably, nearly all the gRNA was in the ∼80S complex regardless of whether it was coexpressed with WT Gag or G2A Gag (Fig 4B, graph). IP with αGFP revealed that both WT and G2A GagGFP were associated with V1B gRNA in the ∼80S putative packaging initiation complex (Fig 4C, graph). Additionally, αGFP IP showed that WT Gag was also associated with V1B gRNA in the ∼500S late assembly intermediate and the ∼750S completed capsids, both of which are formed by WT Gag but not targeting-defective G2A Gag (Fig 4C, graph). While COS-1 cells were used in this experiment to avoid endocytosis of completed virus, as previously described [25], similar results were obtained in human 293T cells expressing WT Gag that were harvested before significant virus endocytosis had occurred (Fig S2A-C). Additionally, coIP results similar to those shown in Fig 4C were obtained even in the absence of PuroHS treatment [25]; thus arguing that PuroHS treatment allows us to study non-translating gRNA without altering the underlying biology. Thus, gRNA is associated with WT Gag in the ∼80S packaging initiation complex, the ∼500S late intermediate, and the ∼750S completed immature capsid. These data are consistent with the ∼80S complex being both the packaging initiation complex and an early assembly intermediate, and with the ∼500S complex being both a late packaging intermediate and late assembly intermediate, as would be expected.

**Fig 4:**
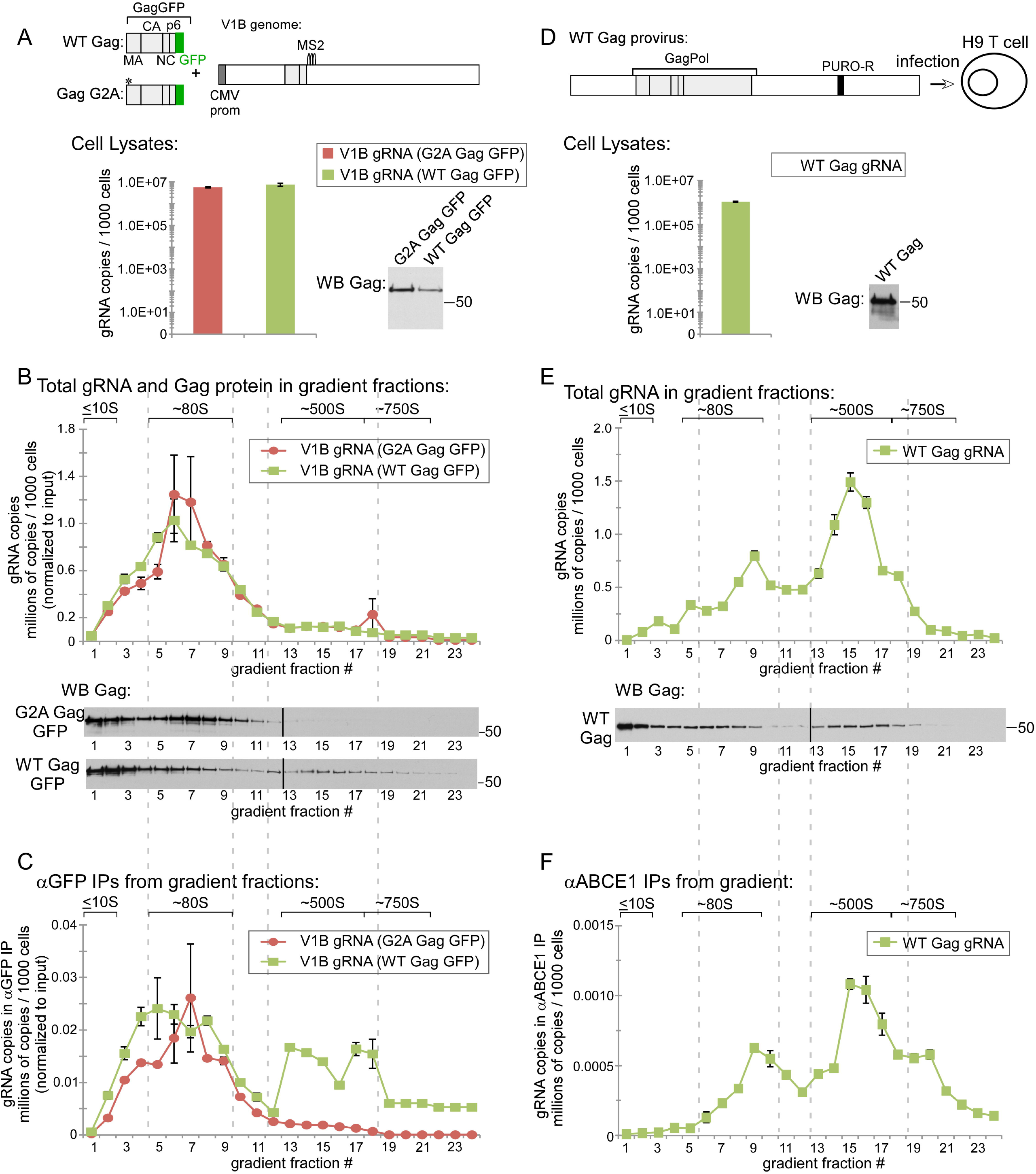
WT Gag forms the packaging initiation complex and a late packaging intermediate, including in infected human T cells. **A)** COS-1 cells transfected with the indicated construct (Fig 1A, Set II) were harvested following PuroHS treatment, and gRNA copy number per 1000 cells was determined. **B)** Lysate from A was analyzed by velocity sedimentation, and gRNA copy number per 1000 cells in each fraction was determined and normalized to the inputs shown in A. **C)** Gradient fractions from B were subjected to IP with αGFP, and gRNA copy number per 1000 cells was determined for IP eluates from each fraction and normalized to input in A. **D)** H9 cells chronically infected with the indicated provirus (H9-HIV; Fig 1A; Set III) were harvested following PuroHS treatment, and gRNA copy number per 1000 cells was determined for each fraction. **E)** Lysate from D was also analyzed by velocity sedimentation, and gRNA copy number per 1000 cells in each fraction was determined. **F)** Gradient fractions from E were subjected to IP with αABCE1 and gRNA copy number per 1000 cells was determined for IP eluates from each fraction. Brackets at top show S value markers, and dotted lines demarcate assembly intermediates based on their WB migrations. Error bars show SEM from duplicate samples. Data in each column are from a single experiment that is representative of three independent repeats.

Thus far, we had shown that 1) both G2A Gag and WT Gag enter an ∼80S RNA granule to form the packaging initiation complex (Fig 3C, 4C; Fig S2C); 2) this ∼80S packaging initiation complex contains the RNA granule protein DDX6 (Fig 3F) and therefore appears to be an RNA granule; and 3) WT Gag also forms a ∼500S packaging/assembly intermediate containing Gag associated with gRNA (Fig 4C; Fig S2C). Additionally, our previous coIP studies had shown that the ∼80S and ∼500S assembly intermediates contain Gag in association with the cellular facilitators of assembly ABCE1 and DDX6 [11,14]. If the packaging initiation complex and late packaging intermediate are also assembly intermediates, we would expect ABCE1 to be associated with gRNA in ∼80S and ∼500S packaging/assembly intermediates formed by WT GagGFP. Indeed, αABCE1 immunoprecipitated V1B gRNA from ∼80S and ∼500S intermediates formed by WT GagGFP (Fig S2D-F), thereby confirming that these two complexes correspond to previously described ∼80S and ∼500S assembly intermediates derived from RNA granules [11,14,25,33]. Thus, gRNA in the ∼80S packaging initiation complex and the ∼500S packaging/assembly intermediate is associated with both WT Gag (Fig 4C, Fig S2C) and ABCE1 (Fig S2F). These data support a model in which Gag enters an RNA granule that contains ABCE1, DDX6, and gRNA to form the packaging initiation complex and remains associated with this granule as it completes packaging and assembly.

### gRNA is present in RNA granule-like packaging intermediates in chronically infected T cells

Previously, we showed that in human H9 T cells chronically infected with HIV (H9-HIV), Gag associates with ABCE1 and DDX6 in ∼80S and ∼500S complexes by coIP [11]. Here we analyzed H9-HIV cells on gradients and found that nearly all the non-nuclear non-translating gRNA is in ∼80S and ∼500S complexes, and that gRNA in those complexes was associated with ABCE1 by coIP (Fig 4D-F). Gag was also present in the ∼80S and ∼500S regions of the gradient, as expected (Fig 4E WB). Interestingly, at steady state in H9-HIV lysate, the ratio of non-translating gRNA in 80S vs. 500S intermediates was shifted towards more ∼500S gRNA, compared to transfected COS-1 or 293T cells (compare 4E, F to 4B,C and S2B, C). Together our findings demonstrate that in three different cell types, including a more physiologically relevant HIV-infected human T cell line, gRNA is only associated with Gag in RNA granule-derived assembly intermediates. Additionally, our findings raise the possibility that formation of the late ∼500S packaging intermediate occurs more efficiently in human T cells.

### Gag requires a gRNA-binding domain to enter the subset of granules containing genomic RNA

Given the longstanding observation that the NC domain is required for association of Gag with gRNA, we would expect NC to be critical for targeting Gag to gRNA-containing RNA granules. To test this possibility, we took advantage of GagZip, in which NC is replaced with a heterologous leucine zipper (LZ). GagZip has been used to demonstrate that NC has two functions during immature capsid assembly - NC binds specifically to gRNA and also promotes oligomerization of Gag via non-specific RNA association [21,24]. Because LZ promotes direct protein-protein interactions, it substitutes for the oligomerization function of NC; thus, GagZip is assembly-competent and produces VLPs [22–24]. However, because LZ does not bind to RNA, these GagZip VLPs lack gRNA or other RNAs [23,24]. Interestingly, we found previously that GagZip forms the ∼80S and ∼500S ABCE1- and DDX6-containing assembly intermediates [11,22], even though it produces VLPs that lack gRNA (Fig 1C). These data raised the possibility that GagZip localizes to a broad class of ABCE1- and DDX6-containing RNA granules, but fails to localize to the subset of these granules that contains gRNA because it lacks the gRNA-binding NC domain.

Before testing this hypothesis, we first confirmed that GagZipGFP, like GagZip, produces VLPs that lack gRNA when cotransfected with the V1B genome *in trans* (Fig S1A and 1C). In cells transfected with WT Gag or GagZipGFP, and V1B *in trans* (Fig 1A, Set II constructs), both Gag proteins were expressed at similar steady state levels as were their gRNAs (Fig 5A), and non-translating V1B gRNA was primarily in the ∼80S RNA granule in both cases (Fig 5B graph). WB confirmed that GagZipGFP forms a prominent ∼500S RNA granule-like assembly intermediate (Fig 5B WB). In addition, previously we confirmed that GagZip also forms the ∼80S RNA granule albeit at lower levels than for WT Gag (in [22], see Fig 4 dark exposures and Fig 5). Notably, αGFP coimmunoprecipitated gRNA from ∼80S and ∼500S fractions of cells expressing WT GagGFP (as in Fig 4C), but αGFP failed to coIP gRNA from any fraction of the GagZipGFP gradient (Fig 5C, graph). Controls showed that αGFP immunoprecipitated GagZipGFP protein as effectively as WT GagGFP from ∼80S and ∼500S fractions (Fig S3A), so the failure to coIP gRNA cannot be attributed to reduced IP efficiency. These data argue that both WT Gag and GagZip localize to ∼80S RNA granules, but WT Gag stably associates with a subset of these RNA granules that contains gRNA, while GagZip does not stably associate with the gRNA-containing subset of these granules. Thus, the NC domain is required to form the packaging initiation complex, and acts by directing Gag to the gRNA-containing granule subset. Moreover, our data suggest that a different region of Gag (present in GagZip but not in MACA) is responsible for bringing Gag to a broader class of RNA granules, most of which lack gRNA.

**Fig 5:**
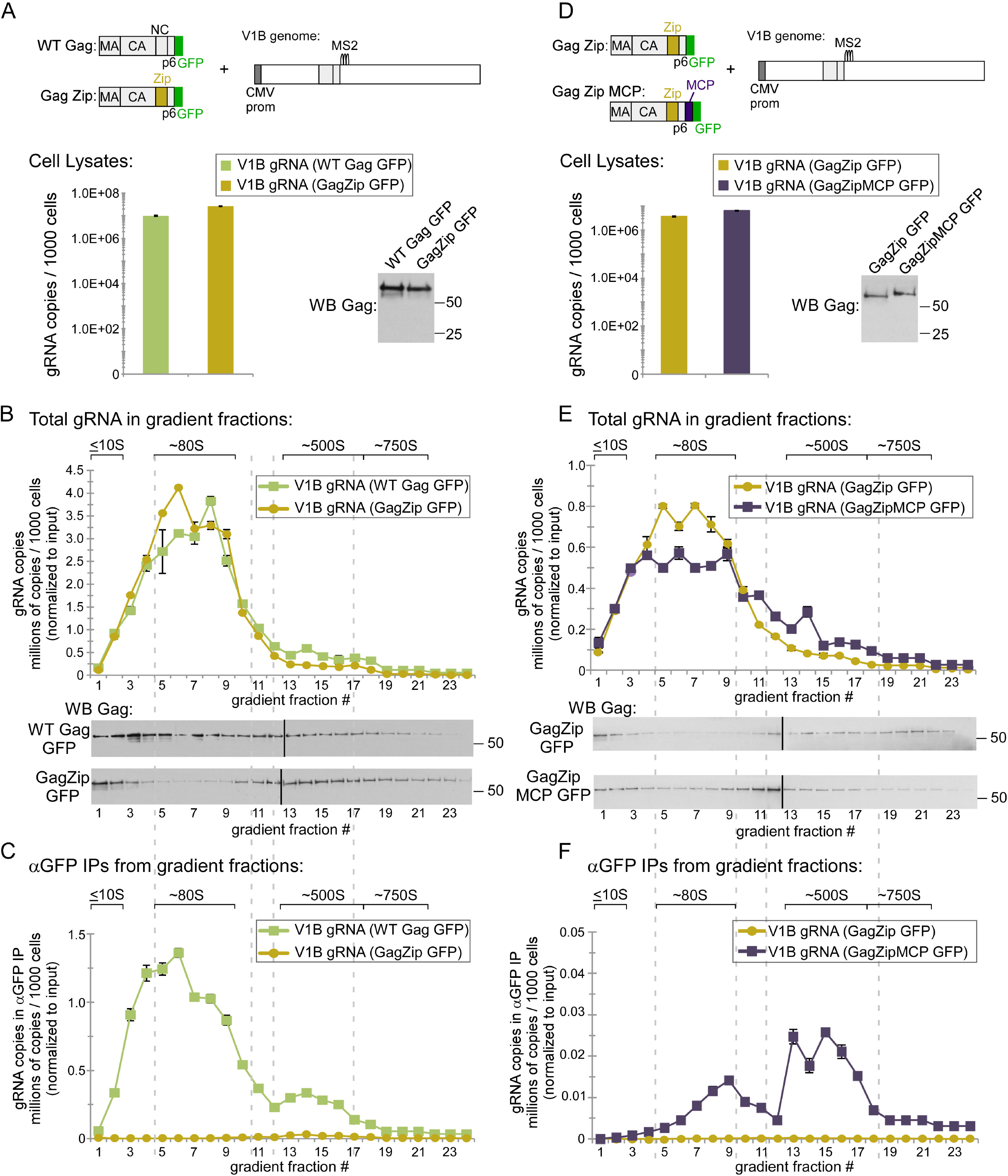
GagZip fails to associate with gRNA-containing granules, but is rescued by a heterologous gRNA-binding domain. **A)** COS-1 cells transfected with indicated constructs (Fig 1A, Set II) were harvested following PuroHS treatment, and gRNA copy number per 1000 cells was determined. **B)** Lysates from A were analyzed by velocity sedimentation, and gRNA copy number per 1000 cells in each fraction was determined and normalized to the inputs shown in A. **C)** Gradient fractions from B were subjected to IP with αGFP, and gRNA copy number per 1000 cells was determined for IP eluates from each fraction and normalized to input in A. **D)** COS-1 cells transfected with indicated constructs (Fig 1A, Set II) were harvested following PuroHS treatment, and gRNA copy number per 1000 cells was determined. **E)** Lysates from D were analyzed by velocity sedimentation, and gRNA copy number per 1000 cells in each fraction was determined and normalized to the inputs shown in A. **F)** Gradient fractions from E were subjected to IP with αGFP, and gRNA copy number per 1000 cells was determined for IP eluates from each fraction. Brackets at top show S value markers, and dotted lines demarcate assembly intermediates based on their WB migrations. Error bars show SEM from duplicate samples. Data in each column are from a single experiment that is representative of two independent repeats.

Finally, we asked whether we could restore targeting of GagZip to a gRNA-containing RNA granule subset via a heterologous gRNA-binding domain. Because the V1B genomic RNA contains MS2 stem loops, we reasoned that the MS2 coat protein (MCP), which binds with high affinity to MS2 stem loops (reviewed in [36]), could serve as a heterologous gRNA-binding domain if inserted into GagZip. Therefore, we generated a construct, here called GagZipMCP, in which MCP is fused to the C-terminus of GagZip. Additionally, we showed that GagZipMCP forms VLPs that contain the V1B genome, unlike GagZip (Fig S3B). When VIB was cotransfected either with GagZip or GagZip MCP, to similar steady state levels (Fig 5D), and analyzed on gradients, gRNA was found largely in ∼80S granules in both cases (Fig 5E). However, αGFP coimmunoprecipitated gRNA from ∼80S and ∼500S fractions formed by GagZip MCP GFP, but not from the corresponding complexes formed by GagZip, analyzed in parallel (Fig 5F). Thus, fusion to MCP, a heterologous gRNA-binding domain, redirected GagZip to the specific subset of RNA granules that contains gRNA, resulting in formation of the gRNA-containing ∼80S packaging initiation complex and ∼500S intermediates. These data confirmed that a gRNA-binding domain is required to target Gag to the subset of granules containing gRNA.

### *In situ* studies confirm that assembling Gag colocalizes with the RNA granule proteins

Next we asked whether we can confirm the presence of Gag in RNA granules in intact cells by visualizing sites where Gag colocalizes with DDX6 or ABCE1 using the proximity ligation assay (PLA). PLA produces fluorescent spots at sites where two proteins are within 40 nm of each other *in situ.* Briefly, this method uses antibodies conjugated to complementary oligonucleotide probes to detect two proteins of interest; the probes anneal only when within the 40 nm range, leading to a rolling circle amplification product that is recognized by a fluorophore-conjugated oligonucleotide [37](Fig 6A). If assembling Gag enters RNA granules containing ABCE1 and DDX6 as indicated by our biochemical studies, then the assembling Gag constructs (WT Gag, G2A Gag, and GagZip) should produce abundant Gag-DDX6 and Gag-ABCE1 PLA spots relative to assembly-incompetent MACA Gag, which fails to enter granules (Fig 3A-C) and does not coIP with DDX6 or ABCE1 [11,25]. To test this, 293T cells were transfected with WT vs. mutant provirus (Fig 1A, Set I) and analyzed for Gag-DDX6 or Gag-ABCE1 colocalization by PLA. Concurrent Gag IF (Fig 6A) allowed us to confirm that the majority of PLA spots were observed in Gag-expressing cells, and to choose fields for quantitation with comparable Gag levels. For cells expressing WT Gag, G2A Gag, or GagZip, quantified fields contained ∼50 Gag-DDX6 PLA spots per cell (Fig 6 B, C), three-fold more than for cells expressing MACA. Similar results were observed for Gag-ABCE1 PLA spots (Fig 7). Some PLA background was expected in cells transfected with MACA provirus, given that DDX6 and ABCE1, like MACA (Fig 3B), are found in the ∼10S fraction and could therefore be in proximity outside of granules. Thus, PLA appears to identify DDX6- and ABCE1-containing RNA granules co-opted by Gag. Moreover, PLA confirms our biochemical studies in which we showed that WT Gag, G2A Gag, and GagZip target to ABCE1- and DDX6-containing RNA granules (Fig 3–5), as well as previous quantitative immunoelectron microscopy studies showing the colocalization of ABCE1 and DDX6 with assembling Gag [11,25,30,33].

**Fig 6:**
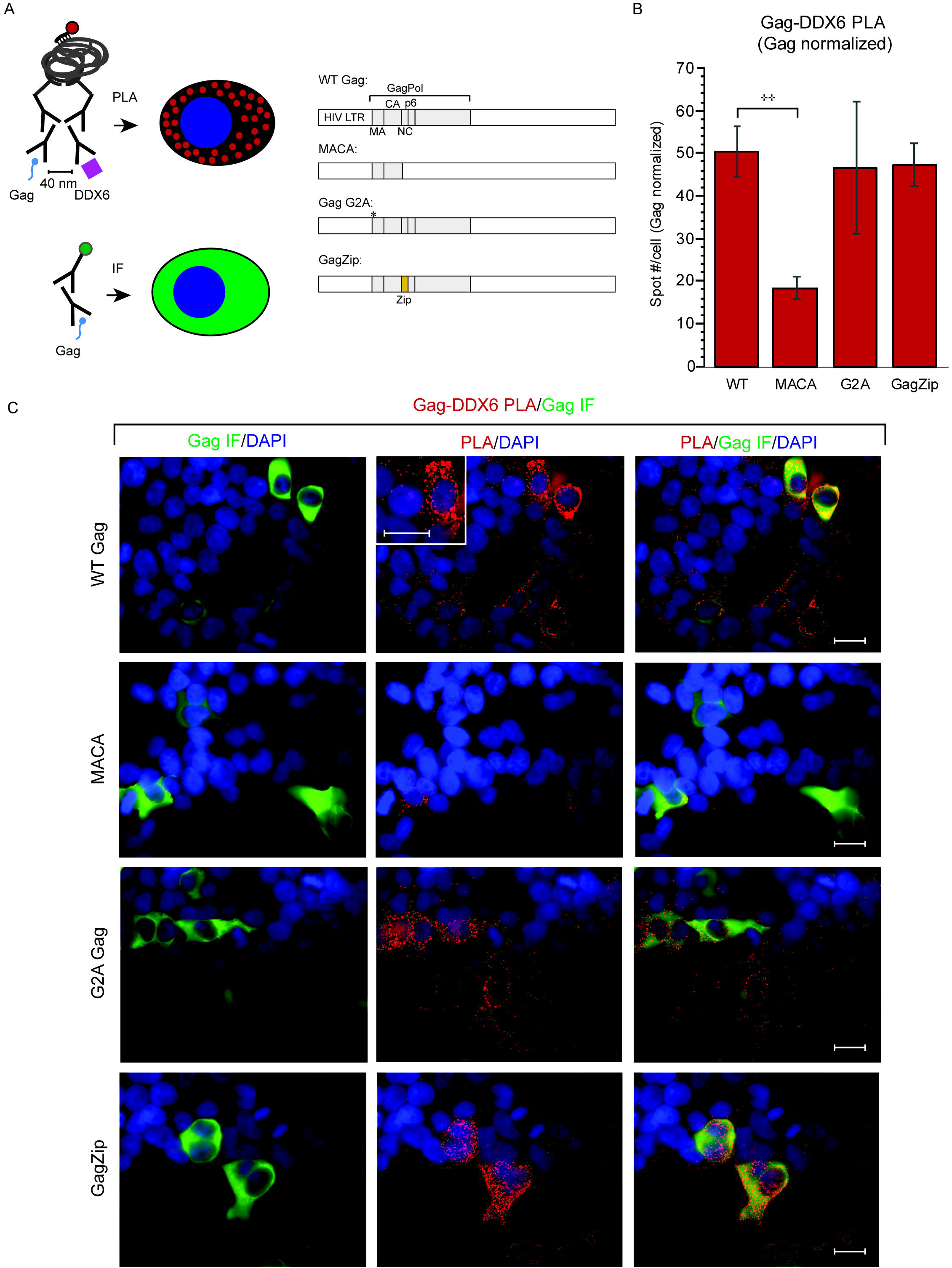
Gag-DDX6 colocalization *in situ* upon provirus expression. PLA was used to detect regions in which Gag is within 40 nm from DDX6 *in situ*, with concurrent αGag IF for quantification of intracellular Gag levels. **A)** Experimental schematic and diagrams of proviruses transfected into 293T cells (Fig 1A, Set 1). **B)** The average number of PLA spots per cell was determined for all Gag-positive cells in five randomly chosen fields, and normalized to Gag levels. Error bars show SEM. **C)** Representative images. From left to right for each construct: Gag IF (green) with DAPI-stained nuclei (blue), Gag-DDX6 PLA signal (red) with DAPI-stained nuclei (blue), and a merge of all three. Merge demonstrates that PLA spots are mainly in Gag-expressing cells. Inset in PLA panel shows a high magnification view of the cell to the right of the inset. Scale bars, 5 µM. Data are representative of three independent repeats. Error bars show SEM (n=10 cells). ** indicates a significant difference relative to WT (p≤0.001).

**Fig 7:**
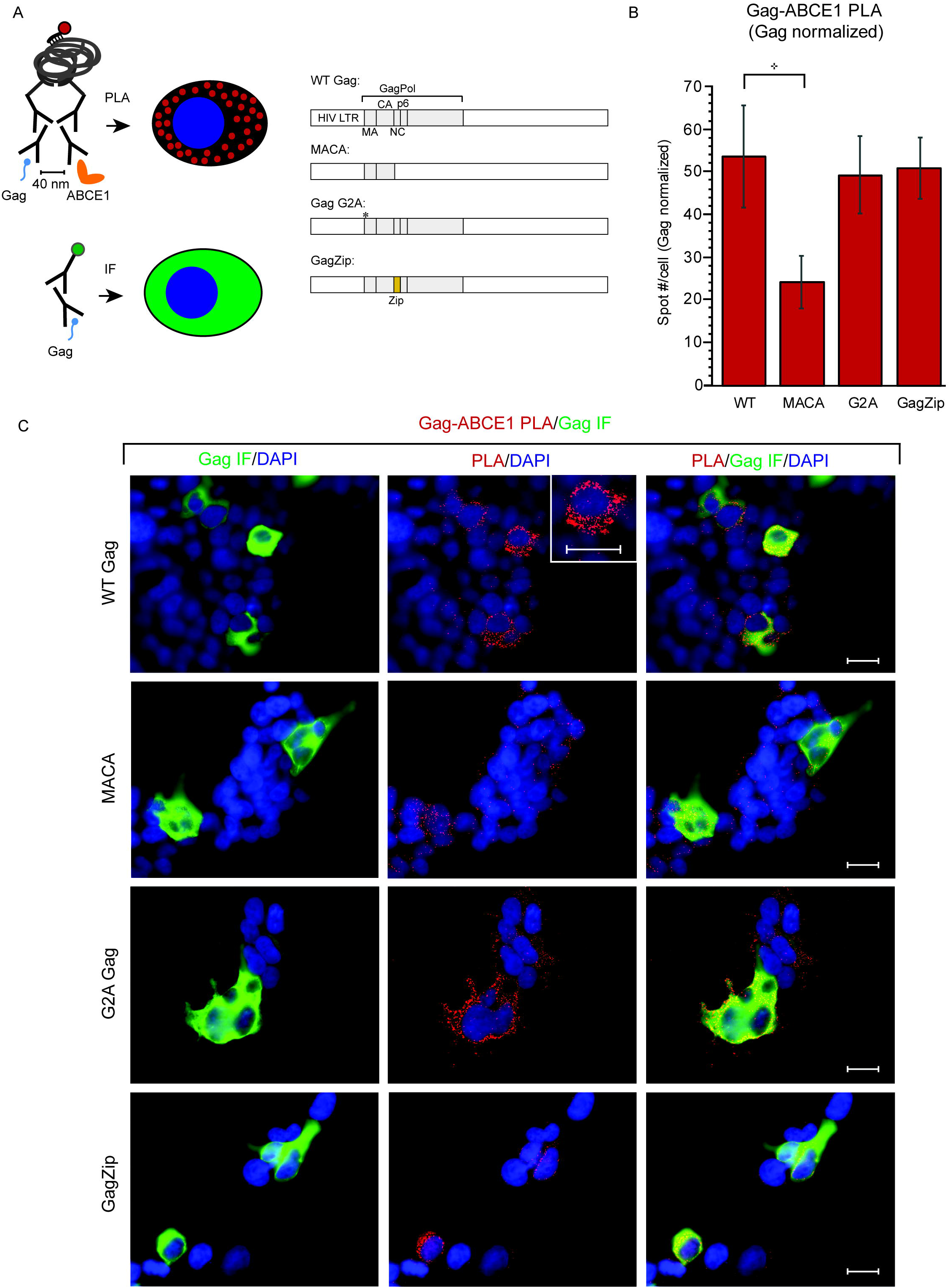
Gag-ABCE1 colocalization *in situ* upon provirus expression. PLA was used to detect regions in which Gag is within 40 nm from ABCE1 *in situ*, with concurrent αGag IF for quantification of intracellular Gag levels. **A)** Experimental schematic and diagrams of proviruses transfected into 293T cells (Fig 1A, Set 1). **B)** The average number of PLA spots per cell was determined for all Gag-positive cells in five randomly chosen fields, and normalized to Gag levels. Error bars show SEM. **C)** Representative images. From left to right for each construct: Gag IF (green) with DAPI-stained nuclei (blue), Gag-ABCE1 PLA signal (red) with DAPI-stained nuclei (blue), and a merge of all three. Merge demonstrates that PLA spots are mainly in Gag-expressing cells. Inset in PLA panel shows a high magnification view of the cell to the left of the inset. Scale bars, 5 µM. Data are representative of three independent repeats. Error bars show SEM (n=10 cells). *indicates a significant difference relative to WT (p<0.01).

### Sites of Gag-DDX6 and Gag-ABCE1 interaction *in situ* are far more numerous than P bodies

While DDX6 is a marker of P bodies, as shown by intense labeling of P bodies by DDX6 IF [31], DDX6 is also found in smaller RNA granules [11]. Given that cells typically contain fewer than ten P bodies [38], our finding that each Z stack image contains ∼50 Gag-DDX6 PLA spots (Fig 6) suggested that granules containing Gag and DDX6 are far more numerous than P bodies. To test this possibility directly, we analyzed 293T cells for G2A Gag-DDX6 PLA spots with concurrent DDX6 IF, to allow detection of PLA spots and P bodies in the same fields (Fig 8A). G2A provirus was used here because

**Fig 8:**
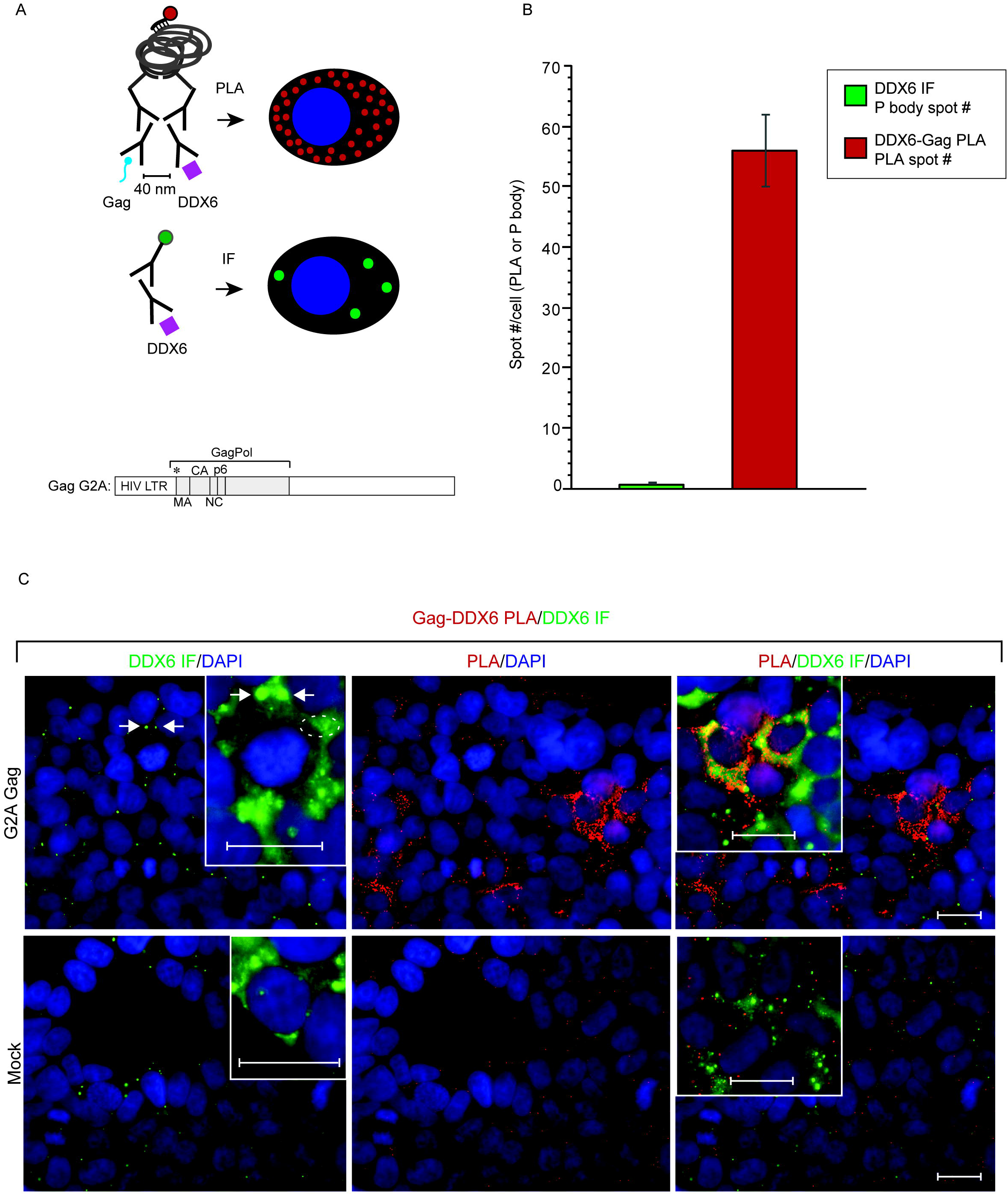
Packaging initiation complexes containing Gag and DDX6 are far more numerous than P bodies

PLA was used to identify regions in which Gag is within 40 nm from DDX6 *in situ*, with concurrent αDDX6 IF for quantification of P bodies. **A)** Experimental schematic. 293T cells were transfected with the indicated G2A provirus (Fig 1A, Set 1) or mock transfected. **B)** The average number of P bodies per cell (green bar) or PLA spots per cell (red bar) was determined for all Gag-positive cells in five randomly chosen fields. Error bars show SEM (n=10 cells). **C)** Representative images. From left to right for G2A Gag and Mock: DDX6 IF to detect P bodies (green) overlaid with nuclei (blue), PLA signal (red) overlaid with nuclei (blue), and a merge of all three. Insets in DDX6 IF panels show a high gain/high magnification version of cell to left, revealing a diffuse, low intensity DDX6 signal (example in dotted oval). Arrows indicate the same two P bodies in low and high magnification images. Insets in merged panels show a high gain/high magnification version of two cells to right. Scale bar, 5 µM. Data are representative of three independent repeats.

G2A Gag is arrested as the DDX6-containing ∼80S packaging initiation complex (Fig 3B, C, F,); thus, most G2A Gag-DDX6 PLA spots likely represent packaging initiation complexes. Each Z stack image from G2A-expressing cells displayed an average of 56 Gag-DDX6 PLA spots, but only one P body by DDX6 IF (Fig 8B, C). Thus, packaging initiation complexes are far more numerous than P bodies. Interestingly, DDX6 IF with high gain revealed a diffuse, low-intensity, granular DDX6 signal (Fig 8C insets), in addition to the bright DDX6 positive P body foci, in both G2A-expressing and mock cells. The low-intensity DDX6 signal could correspond to DDX6-containing ∼80S RNA granules, some of which are co-opted to form packaging initiation complexes.

### Association of gRNA with DDX6 at the PM *in situ* upon expression of WT Gag but not GagZip

Our biochemical studies showed that WT Gag remains associated with the co-opted RNA granule during late stages of assembly (Fig4), suggesting that assembling Gag takes the co-opted gRNA-containing granule to PM sites of budding and assembly. In contrast, we found that GagZip associates with RNA granule proteins at late stages of assembly, but not with the subset of RNA granules that contains gRNA (Fig 5). Thus, we would expect *in situ* approaches to reveal DDX6-containing RNA granules to be present at WT Gag or GagZip PM sites of assembly; however, the granules co-opted by WT Gag should also contain gRNA, while the granules co-opted by GagZip should contain DDX6 but no gRNA. Previously, we used quantitative immunoelectron microscopy (IEM) to demonstrate that RNA granule proteins (ABCE1 and/or DDX6) are recruited to PM sites of WT Gag and GagZip assembly [11,22,33]; however, these studies did not assess whether gRNA was associated with these granules. Here we used quantitative IEM with double labeling for gRNA and DDX6 to ask whether gRNA is associated with RNA granules at PM sites of assembly for WT Gag vs. GagZip *in situ* (Fig 9). HeLa cells stably expressing MCP fused to GFP (HeLa-MCP-GFP cells) were transfected with V1B genomic constructs encoding MS2 binding sites and Gag *in cis* (Fig 9A; Fig 1A, Set IV constructs; phenotypes confirmed in Fig S1B). Sections were labeled with αDDX6 (large gold) to mark RNA granules, and with αGFP to mark MCP-GFP-tagged gRNAs (small gold). PM assembly sites, defined by the presence of membrane deformation consistent with budding, were quantified and scored for gRNA labeling, DDX6 labeling, and double labeling (Fig 9B, C; Table S1). When all sites of DDX6 labeling at PM assembly sites were quantified, similar high levels of DDX6 labeling were observed at both WT and GagZip PM assembly sites (56% vs. 40% of all WT vs. GagZip PM assembly sites displayed DDX6 labeling, respectively, p>0.01; shown as total DDX6+ in Fig 9B; shown as D+ tot in Table S1). Notably, quantitation of all sites of gRNA labeling at the PM revealed that gRNA was significantly more common at WT assembly sites relative to GagZip PM assembly sites (64% vs. 20% of all WT vs. GagZip PM assembly sites displayed gRNA labeling, respectively, p<0.005; shown as total gRNA+ in Fig 9B; shown as g+ Tot in Table S1). Our most striking results were obtained upon quantitation of PM assembly sites that were double labeled for DDX6 and gRNA. Abundant gRNA and DDX6 double labeling was observed at WT PM assembly sites, but not at GagZip PM assembly sites (33% vs. 5% of all WT vs. GagZip PM assembly sites, respectively, displayed double labeling, p<0.005; shown as gRNA+/DDX6+ in Fig 9B; shown as g+D+ in Table S1). As expected, the assembly-incompetent MACA formed very few early or late PM assembly sites, unlike WT and GagZip. The same patterns were observed when early and late assembly sites were analyzed separately (Table S1). Thus, IEM analysis of PM assembly sites supports our conclusion that both WT and GagZip co-opt RNA granules during packaging and assembly, but only the RNA granules co-opted by WT Gag also contain gRNA. Moreover, these quantitative IEM studies (Fig 9) along with our PLA studies (Fig 6–8) provide *in situ* validation, in the absence of PuroHS treatment, of our biochemical studies.

**Fig 9.**
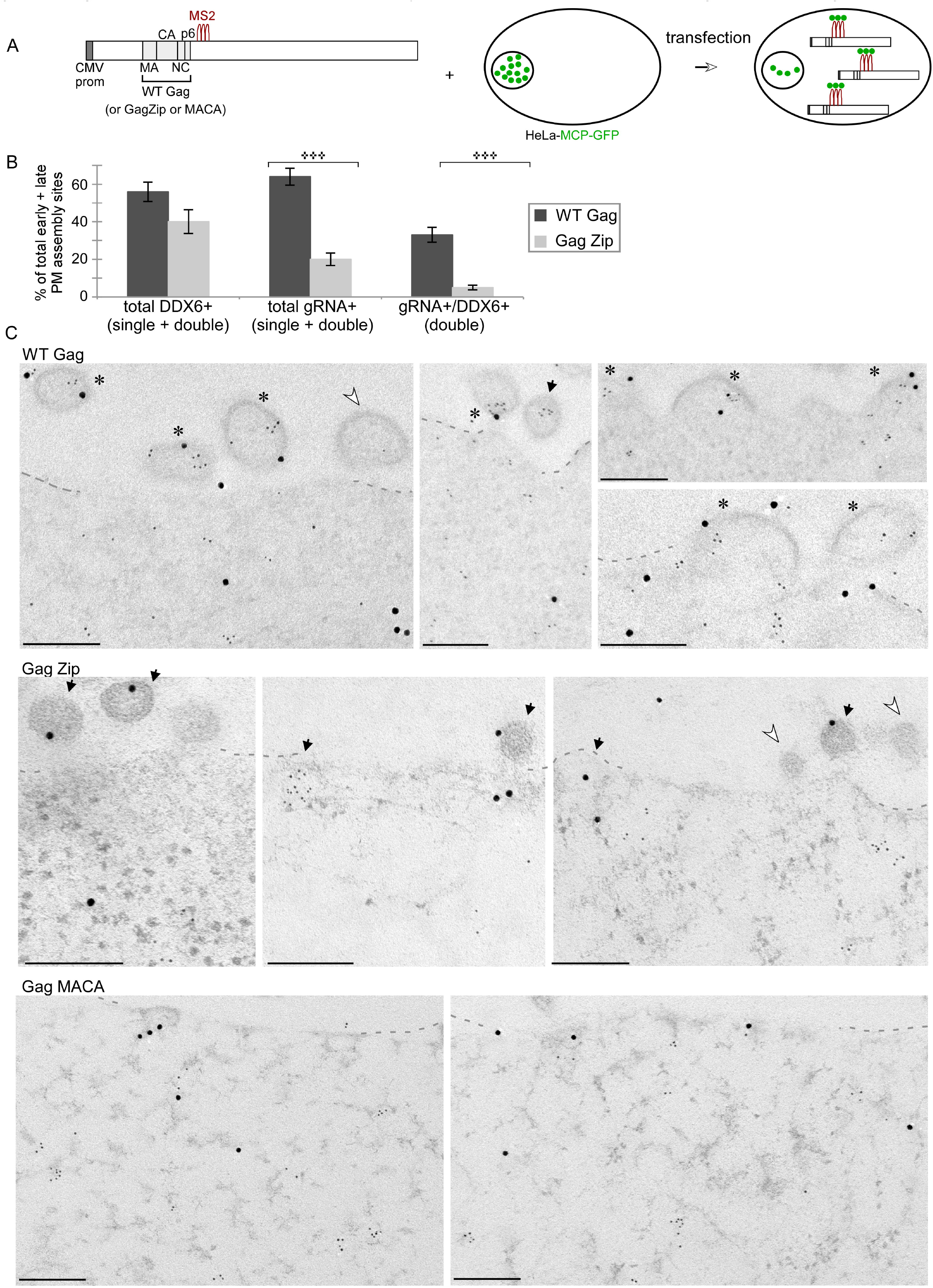
The gRNA-DDX6 association at PM assembly sites is observed *in situ* for WT Gag but not for GagZip. **A)** HeLa cells expressing MCP-GFP were transfected with V1B genomes that contain MS2 binding sites (Fig 1A, Set IV constructs) and express WT Gag (shown), GagZip, or MACA (diagrams not shown). Cells were analyzed by double-label IEM, using αDDX6 (large gold), and αGFP, which allowed detection of MCP-GFP labeled gRNA (small gold). All early and late assembly sites at the PM were identified in ten cells (∼250 µm of PM total per group), and scored for gRNA labeling, DDX6 labeling, and double labeling. **B)** Graph shows the percentage of all early and late PM assembly events that are DDX6-labeled (total DDX6+), gRNA-labeled (total gRNA +), or double-labeled (gRNA+/DDX6+). Error bars show SEM (n=10 cells). *** indicates a significant difference between WT Gag and GagZip (p<0.005). For additional data, see Table S1. C) Images show representative assembly sites at the PM for each group, with symbols indicating early or late PM assembly sites that are single-labeled for either gRNA or DDX6 (dark arrows), double-labeled (asterisks), or unlabeled (open arrows). Dotted lines outline the PM. Scale bars, 200 nm.

## Discussion

It is increasingly evident that, within cells, the fate of an RNA is determined in large part by its associated cellular proteins [39]. For this reason, it will be important to determine whether HIV-1 packaging initiation occurs in the context of cellular RNA binding proteins. Here we showed that almost all of the non-nuclear, non-translating gRNAs in HIV-1 expressing cells are found in large RNPCs (>30S). Moreover, a subclass of these RNPCs, identified as ∼80S host RNA granules, is co-opted to form the packaging initiation complex. We identified this ∼80S host RNA granule as the packaging initiation complex by demonstrating that it is the only gRNA-containing complex formed by the G2A Gag mutant, which arrests at packaging initiation (Fig 3C). We also demonstrated that the RNA-granule-derived ∼80S packaging initiation complex contains the host enzymes ABCE1 and DDX6, and is found in HIV-1-infected human T cells. Additionally, our studies revealed that Gag does not require a gRNA-binding domain to stably associate with ∼80S host RNA granules, but does require a gRNA-binding domain to stably associate with the subset of these granules that contains gRNA. Finally, we showed that packaging initiation complexes are far more numerous than P bodies *in situ*; thus, packaging initiation complexes are not equivalent to P bodies, consistent with our finding that packaging initiation complexes correspond to smaller ∼80S RNA granules.

Based on our findings, we propose the following model for initiation of gRNA packaging (Fig 10). In the cytoplasm, gRNA is either in translating complexes or in non-translating host RNPCs of ∼30S-80S. WT Gag uses a two-step mechanism to co-opt gRNA-containing granules to form the ∼80S packaging initiation complex. One of these two steps involves an event that is independent of gRNA binding, and localizes WT Gag to a broad class of ABCE1- and DDX6-containing ∼80S RNA granules. The second step involves a gRNA-binding-dependent event that localizes WT Gag to the subset of these granules that contains gRNA. In the case of WT Gag, targeting, packaging, and late stages of multimerization continue in association with this RNA granule, leading to formation of the ∼500S packaging/assembly intermediate and the ∼750S completely assembled capsid, both of which contain gRNA. The packaging initiation complex and the ∼500S packaging/assembly intermediate contain ABCE1 and DDX6, two host enzymes that facilitate virus assembly [11,14] and that may distinguish this subclass of granules. In contrast, the intracellular ∼750S completed capsid has dissociated from the RNA granule [11,14,32,33] and subsequently completes budding and release. Three caveats to this model should be noted. First, how gRNA enters the host RNA granules is not known. Most likely, gRNA-containing RNPCs are generated during transcription and undergo successive rounds of remodeling during nuclear export and in the cytoplasm to form ∼80S non-translating RNA granules and translating complexes in the cytoplasm. Secondly, while we refer to complexes in the ∼80S region as a single ∼80S complex for simplicity, future studies will be required to determine if complexes in this region are homogenous or display some heterogeneity. Thirdly, while our coIP studies demonstrate that Gag is associated with gRNA in the packaging intermediates, we do not know whether Gag and gRNA make direct contact with each other in these complexes. Based on studies by others [40], we speculate that Gag proteins make direct contact with only a few regions of gRNA in the ∼80S packaging initiation complex, but contact many more regions of the gRNA in the ∼500S late packaging intermediate.

**Fig 10.**
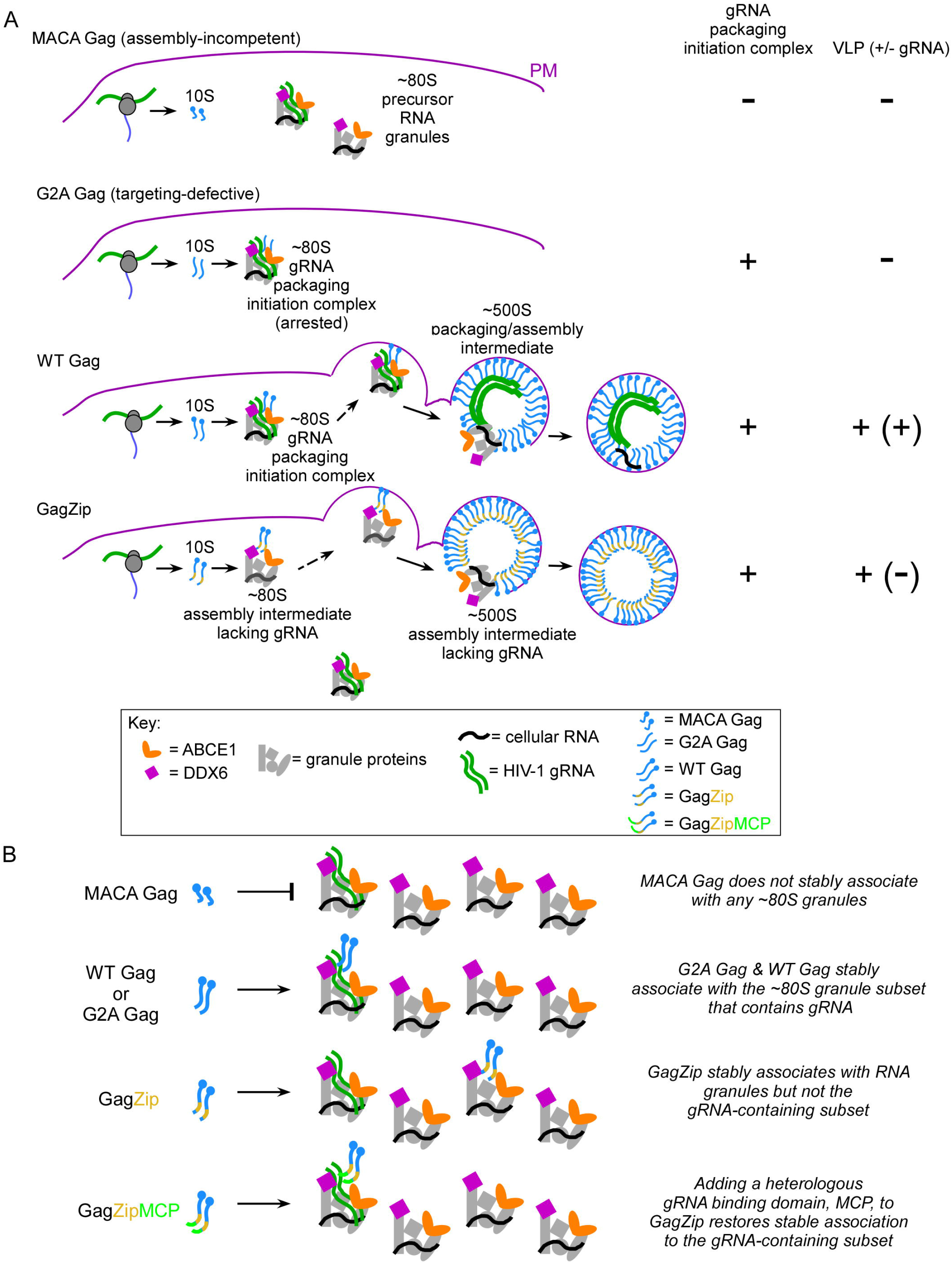
Model for how HIV-1 gRNA packaging is initiated within a subclass of host RNA granules. **A)** MACA Gag, which lacks NC, fails to associate with RNA granules. In contrast, the targeting-defective G2A Gag mutant associates with ∼80S RNA granules that contain gRNA, leading to packaging initiation complex formation in the cytoplasm without further progression. WT Gag also associates with RNA granules and forms the packaging initiation complex in the cytoplasm, which is then targeted to the PM where Gag multimerizes to complete assembly. RNA granule proteins dissociate from the fully assembled capsid prior to VLP budding and release. GagZip also targets to ∼80S RNA granules, but because it contains a protein-protein dimerization domain in place of the gRNA-binding NC domain it is unable to target to the subset of ∼80S granules that contains gRNA. Ultimately, GagZip also undergoes PM targeting, assembly, dissociation of RNA granule proteins, and VLP release; however, GagZip VLPs lack gRNA because the gRNA-binding-deficient GagZip targeted to the “wrong” subset of ∼80S granules. **B)** The two-step model for initial targeting of Gag to gRNA-containing RNa granules is shown in more detail. Only a subset of ∼80S RNA granules contain gRNA. While MACA Gag does not associate with any RNA granule subsets, G2A and WT Gag target to the subset of RNA granules that contains gRNA. GagZip targets to RNA granules regardless of whether they contain gRNA; however, a heterologous gRNA-binding domain (MCP) is able to rescue GagZip, allowing it to target to gRNA-containing ∼80S granules.

A stunning finding of this study is that the packaging initiation complex has an S value similar to the eukaryotic ribosome, a large RNPC containing four RNAs and 79 cellular proteins. Interestingly, studies by others are consistent with this finding. Specifically, the reported diffusion coefficient of ∼70S bacterial ribosomes (0.04 µm^2^ sec^−1^; [41] is similar to the ∼0.07 and 0.014 to µm^2^ sec^−1^ diffusion coefficients reported for cellular subpopulations of HIV-1 gRNA [9] and Gag [10], respectively. Our finding that the packaging initiation complex is ∼80S suggests that this complex contains numerous additional components, since Gag is likely a dimer or small oligomer at this stage [7], and a Gag dimer on its own would be ∼5S. Previously we showed that two other viral proteins are present in assembly intermediates - HIV-1 GagPol [33] and Vif [14]; however, it is likely that host components account for much of the molecular mass of the ∼80S packaging initiation complex. Here we showed that the packaging initiation complex contains at least two host enzymes that are known to facilitate HIV-1 capsid assembly, ABCE1 and DDX6 [11,14]. In addition, our earlier studies demonstrated that the RNA granule proteins AGO2 and DCP2 are also present in these RNA-granule-derived assembly/packaging intermediates [11]. Further studies will be required to define the exact proteome and RNAome of the ∼80S packaging initiation complex and determine whether this complex contains other RNA binding proteins such as Staufen1, which plays a role in HIV-1 packaging and assembly [42–44], and M0V10, which is packaged by HIV-1 and modulates virus infectivity [45,46].

Importantly, using PLA we showed that although the packaging initiation complex contains the P body marker DDX6, it is not equivalent to a P body, and corresponds instead to granules that are much smaller than P bodies (Fig 8). The PLA findings correspond well with our biochemical findings, which suggest that the ∼80S granule should be similar in size to the ∼80S ribosome, which is ∼25 nm in diameter [47]; thus the DDX6-containing ∼80S granule should be four to twelve times smaller than DDX6-containing P bodies, which range from 100 – 300 nm in size [48]. These findings are also consistent with an earlier study showing that Gag and gRNA are not found in P bodies [49]. Note, however, that the co-opted ∼80S RNA granules could represent P body subunits or could exchange with P bodies. Thus, we cannot rule out the possibility that the packaging initiation complex is occasionally found in P bodies, perhaps explaining an earlier report [50]. Future studies will need to compare the components of P bodies with those of the ∼80S RNA granule that contains the vast majority of the gRNA in HIV-1 infected cells. However, it is worth noting that in the case of the Ty3 yeast retrotransposon, which is distantly related to HIV-1, assembling Ty3 Gag and its packaged Ty3 RNA are found in large clusters that contain the yeast DDX6 homologue dhh1 [51,52]; moreover knockdown and mutational analyses indicate a role for RNA granule proteins in formation of functional Ty3 particles in yeast [53,54]. Similarly, we had previously shown using siRNA knockdown that DDX6 promotes immature HIV-1 capsid assembly, and can be rescued by a siRNA resistant WT DDX6 but not a ATPase-defective DDX6 mutant [11]. While others have not observed that DDX6 is required for HIV-1 capsid assembly [49], this could be because other helicases in the co-opted RNA granule can substitute for DDX6 in some cell types or when given enough time to undergo upregulation.

Our identification of the two-step mechanism for stable association of Gag with gRNA-containing ∼80S granules resulted from the observation that an assembling Gag that lacks a gRNA-binding domain (GagZip) targets to RNA granules that lack gRNA, but can be redirected to gRNA-containing granules by addition of a heterologous gRNA-binding domain (Fig 5C, F; [11,22]). Thus, it appears that most ∼80S RNA granules contain cellular RNAs, with only a small subset containing gRNA. Notably, a key difference between MACA, which fails to enter RNA granules, and GagZip, which enters ABCE1- and DDX6-containing granules that lack gRNA, is that GagZip has a heterologous oligomerization domain (LZ) that substitutes for NC, a domain in WT Gag that mediates oligomerization. Together, these data argue that two events are required for stable association of Gag with gRNA-containing granules, with one being the aforementioned poorly understood, NC-independent step in which oligomerization-competent Gag targets to a large class of ∼80S ABCE1- and DDX6-containing RNA granules, most of which lack gRNA; and the other being a gRNA-binding step, dependent on NC or a heterologous gRNA-binding domain, that enables stable association with a gRNA-containing subset of these granules.

Our finding that, in WT Gag, the NC domain is required for packaging initiation complex formation fits with decades of studies showing the importance of NC in packaging. At the same time, our data argue for a new view of packaging. To date, packaging studies have not specified where or how Gag associates with gRNA; we fill this gap by providing evidence that packaging is initiated within a unique host RNPC that contains RNA granule markers and nearly all the packageable gRNA. Interestingly, sequesteration of gRNA within unique host RNA granules could be both beneficial and problematic for the virus. On the one hand, gRNA sequestration is likely to be highly advantageous to the virus - it could allow gRNA to evade detection by the host immune system, create a site where assembling Gag can be concentrated, and provide Gag access to RNA helicases that could facilitate displacement of host RNA binding proteins from gRNA allowing them to be replaced with Gag. On the other hand gRNA sequestration creates a dilemma for the virus in that it puts gRNA in a different subcellular compartment (the RNA granule) than newly translated Gag, which is found either associated with translating ribosomes or in the soluble compartment. To solve this problem, Gag appears to have evolved a mechanism for localizing to a subclass of mRNA-containing host RNA granules, which in turn allows efficient NC-dependent localization to the small subset of these granules that contains gRNA. Future studies will need to further test this model and address how both the cell and the virus utilize this unique class of RNA granules.

## Materials and Methods

### Plasmids and cells

Four types of expression systems were utilized in this study. Proviruses (Set I in Fig 1A) are from the LAI strain and were described previously [22,33,55]. These have native HIV-1 sequences and the HIV-1 LTRs. Proviruses were transfected into COS-1 or 293T cells (both obtained from the American Type Culture Collection (ATCC), Manassas, VA) as indicated for some biochemical studies and all PLA studies. For other biochemical studies, we used an *in trans* expression system in which the genome is provided by V1B, a modified proviral plasmid that expresses an RNA encoding an assembly-incompetent truncated *gag* gene, all cis-acting packaging signals, full-length *tat, rev,* and *vpu* genes, and 24 MS2 stem loops that bind to MCP [8]. V1B was transfected into COS-1 or 293T cells with WT and mutant SynGagGFP constructs (here referred to as GagGFP constructs) provided *in trans,* where indicated. V1B and the codon optimized SynGagGFP WT and G2A constructs (Set II in Fig 1A) were provided by P. Bieniasz (Rockefeller University, New York, N.Y.) and were utilized in previous live imaging studies [8] and coIP studies [7]. Additional GagGFP constructs (MACA and GagZip) were generated from the WT SynGagGFP construct. For G2A, the glycine in position 2 of Gag was converted to an alanine by via site-directed mutagenesis, as described previously [33]; for GagZip, a leucine zipper was inserted in place of nucleocapsid using Gibson assembly, as described previously [22]. For IEM experiments, HeLa cells that express MCP-NLS-GFP as described previously [8] were obtained from P. Bieniasz (Rockefeller University, New York, N.Y.). To ensure that all HeLa-MCP-GFP cells expressed both Gag and genome following transfection, V1B constructs expressing WT and mutant Gag *in cis* were generated by inserting relevant Gag coding regions from HIV-1 proviruses (Set I in Fig 1A) into V1B constructs containing MS2 binding sites via Gibson assembly to generate Set IV constructs in Fig 1A. Oligos used for site-directed mutagenesis and Gibson assemblies are available upon request.

H9 cells (ATCC) were used to generate H9-HIVpro-cells (Set III in Fig 1A) by infection with virus. Plasmid used for virus production was generated by inserting three protease inactivation mutations into a previously described HIV-1 provirus, LAI strain, that encodes deletions in *env* and *vif,* a frameshift in *vpr,* and substitution of *nef* with a puromycin resistance gene, as described previously [11]. 293T cells were transfected with this plasmid to produce virus, and H9 cells were infected with this virus and maintained under puromycin selection as described previously [32].

COS-1 and 293T cells were maintained in DMEM (Life Technologies) with 10% FBS. H9 cells were maintained in RPMI (Life Technologies) with 10% FBS under puromycin selection. HeLa-MCP-GFP cells were obtained from P. Bieniasz and were maintained in DMEM with 10% FBS, and periodically subjected to blastocidin selection.

### Transfection, IP, and WB

COS-1, 293T, or HeLa-MCP-GFP cells were transfected with 1-5µg DNA using polyethylenimine (Polysciences, Warrington, PA). Cell lysates were harvested in 1X Mg^+2^-containing NP40 buffer (10 mM Tris-HCl, pH 7.9, 100 mM NaCl, 50 mM KCl, 1 mM MgCl, 0.625% NP40) in the presence of freshly prepared protease inhibitor cocktail (Sigma, St Louis, MO) and RNaseOUT (Invitrogen). Where indicated, lysates were treated with 1 mM puromycin HCl (Invitrogen) for 10 min at 26°C followed by 0.5M KCl for 10 min at 26 °C. Lysates were analyzed by WB or RT-qPCR, or subjected to IP as described below. Alternatively, lysates were analyzed by velocity sedimentation, as described below, and gradient fractions were then analyzed by WB, RT-qPCR, or IP.

Except where indicated, lysates or gradient fractions were subjected to IP with affinity purified antibody to ABCE1 [14], HIV immunoglobulin NIH AIDS Reagents Catalog #3957, from NABI and NHLBI), a monoclonal to GFP (Roche), or an antibody to DDX6 (#461, Bethyl Laboratories, Montgomery, TX), using protein G-coupled Dynabeads (Life Technologies). IP eluates were analyzed by SDS-PAGE, followed by western blot (WB) using the primary antibodies described above or a monoclonal antibody directed against HIV-1 Gag p24 (HIV-1 hybridoma 183-H12-5C obtained from Bruce Chesebro through the AIDS Reagent Program Division of AIDS, NIAID, NIH), followed by an HRP-conjugated anti-human IgG secondary antibody (Bethyl Laboratories, Montgomery, TX), or an HRP-conjugated anti-mouse-IgG_1_ (Bethyl Laboratories) or anti-mouse-IgG or anti-rabbit secondary antibody (Santa Cruz Biotechnology, Dallas, TX). In Suppl. Fig 3A, IPs were subjected to WB with HIV immune globulin (provided by NABI and NHLBI, catalog no. 3957 in the AIDS Reagent Program Division of AIDS, NIAID, NIH) for detection of Gag. WB signals from IP eluates were detected using Pierce ECL substrate (Thermo Fisher Scientific) with Carestream Kodak Biomax Light film. For detection of Gag in total cell lysates, velocity sedimentation fractions, and membrane flotation fractions, WBs were performed as described above, or using antibodies conjugated to infrared dyes (LI-COR Biosciences, Lincoln, NE). Quantification of Gag bands on film was performed using Image J software or LI-COR Odyssey software.

### RNA quantification

Transfected cells or VLPs were harvested as described above. Where indicated lysates were analyzed by velocity sedimentation and or IP, as described above except that IPs for gRNA quantification were washed four times in detergent buffer and once in non-detergent buffer. For total cell lysate analysis, aliquots corresponding to ∼5 x10^3^ COS-1 cells, 2 x10^4^ 293T cells and 8.5 x10^3^ HeLa-MCP-GFP cells were used. For VLP analysis, aliquots corresponding to VLPs from 1 x 10^5^ 293T cells and 1.5 x10^4^ HeLa-MCP-GFP cells were used. For gradient analysis, ∼4 x10^5^ COS-1 cells, 2 x 10^6^ 203T cells, or 6 x 10^5^ H9 cells were analyzed on a single 5 ml gradient. Aliquots of gradient fractions or gradient IP fractions were treated with proteinase K (Sigma) at a final concentration of 150 µg/ml in 0.1% SDS, followed by total RNA extraction by using Trizol (Ambion). RNA was precipitated with isopropanol, extracted with BCP (Molecular Research Center), pelleted at 12,000xg for 15 min at 4°C, and the RNA pellet was subjected to DNase I (Invitrogen) treatment (2 u per 50 ul reaction). The iScript Advanced cDNA synthesis kit (Bio-Rad) was used to generate cDNA from 10% of the RNA using random hexamer primers at 42°C for 30 min, followed by heat inactivation. An aliquot of cDNA (2.9%) was used for qPCR using SYBR Green (Bio-Rad) to determine RNA copy number. For HIV-1 gRNA qPCR, we used the following oligos that target bp 162 to 269 within the Gag open reading frame in HIV-1 LAI (at the end of MA and start of CA) and result in a 108 bp amplicon: 5’-AGAAGGCTGTAGACAAATACTGGG-3’ (forward); 5’- TGATGCACACAATAGAGGGTTG-3’ (reverse). These oligos detect the full length HIV-1 provirus as well as the V1B genomic construct, but do not recognize the GagGFP constructs that were transfected *in trans,* as expected since the GagGFP constructs were codon-optimized. To determine the copy number of other RNAs, we used the following qPCR oligos: Tat mRNA, 5’-TCT ATC AAA GCA ACC CAC CTC-3’ (forward) and 5’- CGT CCC AGA TAA GTG CTA AGG-3’ (reverse); 28S rRNA, 5’-CCC AGT GCT CTG AAT GTC AA-3’ (forward) and 5’-AGT GGG AAT CTC GTT CAT CC-3’ (reverse); GAPDH mRNA, 5’-AGG TCA TCC CTG AGC TGA AC-3’ (forward) and 5’-GCA ATG CCA GCC CCA GCG TC-3’ (reverse); and 7SL RNA 5’-GCT ATG CCG ATC GGG TGT CCG-3’ (forward) and 5’-TGC AGT GGC TAT TCA CAG GCG-3’(reverse). All qPCR samples were analyzed in duplicate using the MyiQ RT-PCR detection system and iQ5 software (Bio-Rad). Amplicons corresponding to region amplified by qPCR were used to generate standard curves. Duplicate nine-point standard curves were included on every qPCR plate, ranging from 10^1^ copies to 10^8^ copies, with a typical efficiency of ∼90% or greater and an R^2^ of 0.99. Standard curves were able to detect 10 copies per reaction but not 1 copy, thereby setting the detection threshold at 10 copies per reaction, which was equivalent to ∼1000 copies per 1000 cells for inputs and total gRNA from gradient fractions, ∼100 copies per 1000 cells for IPs from total cell lysates or gradient fractions, and ∼50 copies per 1000 cells for VLPs. Minus RNA controls were included in each experiment and were always zero. RT minus controls were also included in each experiment and ranged from 0 - 100 copies per reaction. Mock transfected VLP controls were used to set the baseline in graphs and ranged from 1 - 1000 copies per 1000 cells. For IPs, nonimmune gRNA copy number was analyzed in parallel and was typically 1-2 logs lower than IPs.

Quantification of the total cell number used in each experiment allowed us to represent all qPCR data as number of gRNA copies per 1000 cells, except for Fig 2 data which is presented as % of total gRNA to allow comparison across different RNAs. Log scales were used to display all VLP, which exhibit large differences. Linear scales were used for IP data, which exhibit smaller differences. Note that differences in transfection efficiency resulted in a range of total gRNA copies per 1000 cells between experiments; likewise, IP efficiency also varied between experiments. Thus the exact number of gRNA copies immunoprecipitated in fractions from different experiments varied considerably, but the pattern did not.

### Analysis of VLP production

Supernatants of COS-1 or HeLa-MCP-GFP cells, transfected as described above, were centrifuged at 2000 rpm (910 x g) for 10 min at 4°C, filtered (0.45 µm) to remove remaining cells, and purified through a 30% sucrose cushion in an SW60Ti rotor at 60,000 rpm (370,000 x g) for 30 min at 4°C, as described previously [22].

### Velocity Sedimentation

Transfected COS-1 cells or 293T cells were harvested at 36 h or 15 h post-transfection, respectively, as described above and diluted into 1X NP40 buffer (10 mM Tris acetate pH 7.4, 50 mM KCl, 100 mM NaCl, 0.625% NP-40). For each sample, 120 µl lysate was layered on a step gradient. To resolve complexes of ∼10S to ∼150S (Fig 2, 3), step gradients were prepared from 5%, 10% 15%, 20%, 25%, and 30% sucrose in NP40 buffer without MgCl (10 mM Tris-HCl, pH 7.9, 100 mM NaCl, 50 mM KCl, 0.625% NP40) and subjected to velocity sedimentation in a 5 ml Beckman MLS50 rotor at 45,000 rpm (162,500 x g) for 90 min at 4°C. To resolve from ∼10S to ∼750S, step gradients were prepared from 10%, 15%, 40%, 50%, 60%, 70%, and 80% sucrose in NP40 buffer without MgCl, and subjected to velocity sedimentation in a 5 ml Beckman MLS50 rotor at 45,000 rpm (162,500 x g), for 45 min at 4°C. Gradients were fractioned from top to bottom, and aliquots were analyzed by WB, IP, and/or RT-qPCR as described above. S values were determined, using a published equation[56] and S value markers, as described previously [34].

### Proximity Ligation Assay

293T cells were plated into 6-well dishes containing coverslips with Grace Biolabs CultureWell silicone chambers (Sigma-Aldrich) attached to create four chambers on each coverslip. Cells were transfected with 3 µg of plasmid per well and 16.5 hours later were fixed for 15 minutes in 4% paraformaldehyde in PBS pH 7.4, permeabilized in 0.5% saponin in PBS, pH 7.4 for 10 minutes, and blocked in Duolink blocking solution (Sigma-Aldrich) at 37°C for 30 min. Cells were incubated in primary antibody (described under IP methods above), followed by Duolink reagents (Sigma-Aldrich): oligo-linked secondary antibody, ligation mix, and red or green amplification/detection mix, with washes in between, as per the Duolink protocol. For concurrent IF, cells were incubated for 15 minutes at RT with 1:1000 Alexafluor 594 anti-mouse or Alexafluor 488 anti-rabbit secondary antibody following the final PLA washes. Cover slips were mounted using Duolink In Situ Mounting Media with DAPI, sealed to the glass slides with clear nail polish, allowed to dry for 24 h at RT, and stored at −20°C. Imaging was performed with a Zeiss Axiovert 200M deconvolution microscope using Zeiss Plan-Apochromat 63X/ aperture 1.4 objective with oil immersion, with AxioVision Rel. 4.8 software. For quantification, five fields containing at least three IF or PLA positive cells were chosen at random and imaged using identical exposure times for the red and green channels (red/green exposures were 1 second/1.5 seconds for Figure 6, 2 seconds/1 second for Figure 7, and 40 milliseconds/250 milliseconds for Figure 8). Images were captured as ten 1-µm Z-stacks centered on the focal point for the PLA. Images were deconvolved using the AxioVision software, then exported as .tif files, and Image J was used to outline Gag-positive cells in each field. Within those positive cells, the central Z-stack image was used to count PLA “spots”, and quantify IF intensity where indicated, using Image J. PLA spot number for each field was then normalized to the average IF intensity within that field, and the results were plotted with error bars representing the SEM for five fields. For Figure 8, Gag-DDX6 PLA was done with either Gag IF or DDX6 IF, and the PLA paired with Gag IF was used for PLA spot quantitation to exclude background spots in Gag-negative cells, whereas the PLA paired with DDX6 IF was used for P body quantitation. For the sample field images used in each figure, after imaging, the red channel gain was increased proportionally (to 3 in Figure 6, to 11 in Figure 7, and to 7 in Figure 8) in the AxioVision Rel. 4.8 software for all conditions, to allow better display of red spots in final figures. The same was done for the green channel gain (increased to 7) to display smaller DDX6 granules in the Fig 8D insets. Images were imported in 8-bit color into Adobe Illustrator to create the final figure layout, without further adjustments to color balance or gamma correction.

### Quantitative IEM

HeLa-MCP-GFP cells were transfected with the indicated constructs. Cells were harvested at 24 h post-transfection in fixative (3% paraformaldehyde, 0.025% glutaraldehyde in 0.1 M phosphate buffer, pH 7.4), pelleted, and subjected to high pressure freezing using the Leica EMPACT2, followed by freeze substitution. Samples were infiltrated overnight with LR White embedding resin (London Resin Company Ltd, Reading, Berkshire, England) in ethanol, changed to straight LR White, embedded in gelatin capsules (Electron Microscopy Sciences (EMS), Hatfield, PA, USA), and cured overnight in a UV light cryo-chamber at 4^°^ C. Sections (∼50 nm) were placed on grids, treated with 0.05 M glycine for 20 min at RT, rinsed in PBS, blocked for 45 min with 1% bovine serum albumin (EMS), and washed in PBS with 0.1% bovine serum albumin-C (BSA-C) (EMS). For immunogold double-labeling, a previously described peptide-specific antiserum directed against DDX6 was affinity purified, desalted, and concentrated [11]. Grids were blocked in 0.5% BSA-C, then incubated with primary antibody to DDX6 (0.1 mg/ml in 0.5% BSA-C), followed by goat a-rabbit F(ab’)2 fragment secondary antibody conjugated to 15nm gold particles (EMS), with washes after each step. Grids were then labeled with the second primary, mouse antibody to GFP (Roche) at 0.2 mg/ml in 0.1% BSAC with 0.002% Tween, followed by goat a-mouse F(ab’)2 fragment conjugated to 6 nm gold particles (EMS). Fixation, negative staining, imaging with the JEOL-1400 transmission electron microscope, and image acquisition have been described previously [11].

For quantification, images were acquired for ten cells from each of the three groups, with the goal being to analyze similar total PM lengths in each group. Cells were chosen randomly, but excluded for the WT and GagZip groups if they had fewer than ten particles at the PM visible at low power. Images encompassed the area of each cell that contained PM assembly sites, with images obtained for ∼250 µm of PM total per group. The number of assembly sites analyzed within this ∼250 µm of PM are not equivalent since the number of assembly sites depends on VLP phenotype and kinetics. A total of 760 WT events and 409 GagZip events were analyzed, but are shown as number of sites per 25 µm PM per cell in Table 1. Each PM assembly site was scored as genome positive (g+), DDX6+ (D+), or double-labeled (g+D+). The following definitions were used for image analysis: early PM assembly sites were defined as displaying curvature at the membrane but with < 50% of a complete bud; late assembly sites” at the PM were defined as displaying curvature but with ≥ 50% of a complete bud. If early or late sites contained two or more small gold particles within the full circle defined by the bud, they were scored as g+. If these sites contained one or more large gold particles within a 150 nm perimeter outside the full circle defined by the bud (roughly the size of an RNA granule plus space to account for the antibodies and gold particle bound to an antigen at the periphery of such a granule), then they were scored as D+. Table 1 shows the average number of early, late, and early+late PM assembly events per 25 µm of PM per cell (n=10 cells +/- SEM), along with the breakdown of how many of these events were g+ (total vs. single-labeled), D+ (total vs. single-labeled), or g+D+ (double-labeled). In italics are g+, D+, and g+D+ per 25 µm of PM per cell as a percentage of the total for each group. Significance was determined on percentage data using a two tailed t-test; not significant was defined as p > 0.01. Labeling of early+late events as a percent of total early+late events is also shown in graphical form in Fig 8B. As described previously [22], the sensitivity of IEM for capturing colocalization is limited by a number of factors including the fact that the 50 nm sections only capture ≤ 50% of a single capsid, which has a diameter of ∼100-150 nm.

## Acknowledgments

We thank B. Schneider, S. MacFarlane, S. Knecht, and the Fred Hutchinson Cancer Research Center Electron Microscopy Resource for assistance with IEM; and C. Geary and C. W. Peterson for assistance with generating reagents. We thank P. Bieniasz for V1B, SynGag, and MS2 coat protein plasmids, and for the Hela-MCP-GFP cell line. The following reagents were obtained from the NIH AIDS Reagent Program, Division of AIDS, NIAID, NIH: catalog no. 1513 HIV-1 Gag p24 hybridoma (183-H12-5C) obtained from B. Chesebro; and catalog no. 3957 HIV Ig from NABI and NHLBI. We thank M. Emerman, A. Sharma, N. Westergreen, D. Ressler, and V. Swain for comments on the manuscript.

---

**Fig. S1: Packaging initiation and VLP phenotypes for V1B *in trans* and *in cis* constructs**.

**A)** COS-1 cells were cotransfected with WT or mutant codon-optimized Gag fused to GFP (GagGFP) along with a genomic construct (V1B) provided *in trans* (Fig. 1A, Set II). Diagram shows schematic of constructs. Top: Gag WB of cells and VLPs. Graph shows gRNA copy number in VLP pellets from the equivalent of 1000 cells, as determined by RT-qPCR. Bottom: Lysates of cells transfected as in A were subjected to IP with αGFP or nonimmune (N) antibody followed by Gag WB (left), with IP inputs shown (center).

**B)** HeLa cells expressing MCP-GFP were transfected with V1B genomes that contain MS2 binding sites and express WT Gag, G2A, GagZip, or MACA (Fig. 1A, Set IV constructs). Diagram shows schematic of constructs. Also shown is a Gag WB of cells and VLPs. Graph shows gRNA copy number in VLP pellets from the equivalent of 1000 cells, as determined by RT-qPCR, with mock background for VLPs subtracted.

**Fig. S2: Association of gRNA with Gag and ABCE1 in the ∼80S packaging initiation complex and ∼500S late packaging intermediate**.

**A)** 293T cells transfected with the indicated WT Gag construct (Fig. 1A, Set II) were harvested + PuroHS, and gRNA copy number per 1000 cells was determined. **B)** Lysates from B were also analyzed by velocity sedimentation, and gRNA copy number from the equivalent of 1000 cells determined. **C)** Gradient fractions from B were subjected to IP with αGFP, and gRNA copy number in IP eluates per 1000 cells determined. **D)** COS-1 cells transfected with the indicated construct (Fig. 1A, Set II) were harvested and analyzed as in A. **E)** Lysate from D was analyzed as in B. **C)** Gradient fractions from E were subjected to IP with αABCE1, and gRNA copy number in IP eluates from the equivalent of 1000 cells determined. Brackets at top show S value markers, and dotted lines demarcate assembly intermediates based on their migrations in the Gag WB. Error bars, SEM from duplicate samples. Data represent three independent repeats.

**Fig. S3: Immunoprecipitation and VLP analysis of GagZip constructs**

**A)** COS-1 cells were cotransfected with a codon-optimized construct expressing either WT Gag or GagZip fused to GFP and a genomic construct (V1B) provided *in trans.* Cells were harvested following PuroHS treatment and analyzed by velocity sedimentation. Paired gradient fractions were subjected to αGFP IP, followed by WB with HIV immune globulin to allow detection of Gag. **B)** COS-1 cells transfected with indicated constructs (Fig. 1A, Set II) and lysates and VLPs were harvested. WB shows Gag in cell lysates and VLPs. Graph shows gRNA copy number, as determined by RT-qPCR, in VLPs or cell lysates representing the equivalent of 1000 cells.

